# Adaptive spectroscopic visible-light optical coherence tomography for human retinal oximetry

**DOI:** 10.1101/2021.05.28.446197

**Authors:** Ian Rubinoff, Roman V. Kuranov, Zeinab Ghassabi, Yuanbo Wang, Lisa Beckmann, David A. Miller, Behnam Tayebi, Gadi Wollstein, Hiroshi Ishikawa, Joel S. Schuman, Hao F. Zhang

## Abstract

Alterations in the retinal oxygen saturation (sO_2_) and oxygen consumption are associated with nearly all blinding diseases. A technology that can accurately measure retinal sO_2_ has the potential to improve ophthalmology care significantly. Recently, visible-light optical coherence tomography (vis-OCT) showed great promise for noninvasive, depth-resolved measurement of retinal sO_2_ as well as ultra-high resolution anatomical imaging. We discovered that spectral contaminants (SC), if not correctly removed, could lead to incorrect vis-OCT sO_2_ measurements. There are two main types of SCs associated with vis-OCT systems and eye conditions, respectively. Their negative influence on sO_2_ accuracy is amplified in human eyes due to stringent laser power requirements, eye motions, and varying eye anatomies. We developed an adaptive spectroscopic vis-OCT (Ads-vis-OCT) method to iteratively remove both types of SCs. We validated Ads-vis-OCT in *ex vivo* bovine blood samples against a blood-gas analyzer. We further validated Ads-vis-OCT in 125 unique retinal vessels from 18 healthy subjects against pulse-oximeter readings, setting the stage for clinical adoption of vis-OCT.

## Introduction

Visual processing is one of the most oxygen-demanding functions in the human body (1, 2). Diseases such as diabetic retinopathy and glaucoma can compromise visual processing in the retina, leading to irreversible vision loss (2, 3). In response to pathological damages, the retina regulates oxygen supply and extraction to satisfy new metabolic demands (2-8). Therefore, change in oxygen saturation (sO_2_) has been broadly agreed to be a sensitive biomarker for various retinal diseases and may be evident before irreversible vision loss occurs (9, 10). Clinical measurement of sO_2_ shall open a critical window for timely intervention to prevent, slow down, or even reverse disease progressions in patients. However, no existing technology has satisfied the clinical need to measure human retinal sO_2_ noninvasively, accurately, and reproducibly.

Optical coherence tomography (OCT), commonly operating within the near-infrared (NIR) spectral range between 800 nm and 1300 nm, enabled noninvasive retinal imaging at a spatial resolution of a few micrometers (11-13). It has become the “Gold standard” for examining structural damages or therapeutic recoveries in nearly all vision-threatening diseases. However, the low spectroscopic optical contrast in blood within the NIR spectral range confounded OCT’s sO_2_ measurements (14-16).

The recently developed visible-light OCT (vis-OCT) (5, 17-22) has shown great promise in overcoming the contrast limit since visible light is highly sensitive to the optical absorption and scattering spectral signatures of blood (14, 23). Vis-OCT analyzes the improved optical spectral contrasts in the blood to measure sO_2_ using short-time Fourier transforms (STFTs) and inverse least-squares fit regression (17). Operating between 510 nm – 610 nm, vis-OCT can provide 1.4-µm full-bandwidth axial resolution (24) and 9-µm STFT axial resolution in the retina. Therefore, vis-OCT can specifically isolate the attenuation spectrum of blood inside individual retinal vessels. This capability provides significant advantages in accuracy over fundus-photography-based retinal oximetry (25), which is not depth-resolved and failed to isolate the attenuation spectrum of blood.

However, reported vis-OCT oximetry techniques (17, 22, 26) are unsatisfactory, especially in human imaging, which hampered its clinical impact. More specifically, current vis-OCT oximetry failed to remove key spectral contaminants (SC), which we found to impact sO_2_ accuracy significantly. Here, we define SC as any erroneous spectra not associated with blood attenuation. We classify SCs into two categories: sample-dependent and system-dependent. When SCs are not correctly accounted for, they contribute to errors in the STFT spectrum and lead to incorrect sO_2_ measurement after inverse least-squares fit.

Fig. 1 is an illustration of the human retina and sources of sample-dependent SCs. The spectral signatures of vis-OCT signals contain contributions from three groups of detected photons: specularly-reflected photons (highlighted by group 1), backscattered photons without interacting with blood (highlighted by group 2), and backscattered photons with interaction with blood (highlighted by group 3). Since the vis-OCT signal is depth-resolved, photon interactions with tissue beneath a blood vessel do not add additional SC. However, photon interactions with tissues above a vessel, such as inner limiting membrane (ILM), retinal nerve fiber layer (RNFL), or vessel wall and red blood cell (RBC), add SC to sO_2_ measurements. To accurately measure sO_2_, vis-OCT needs to separate depth-resolved optical absorption and scattering spectral signatures of the blood from SCs contributed by these three groups of photons.

**Figure 1.**
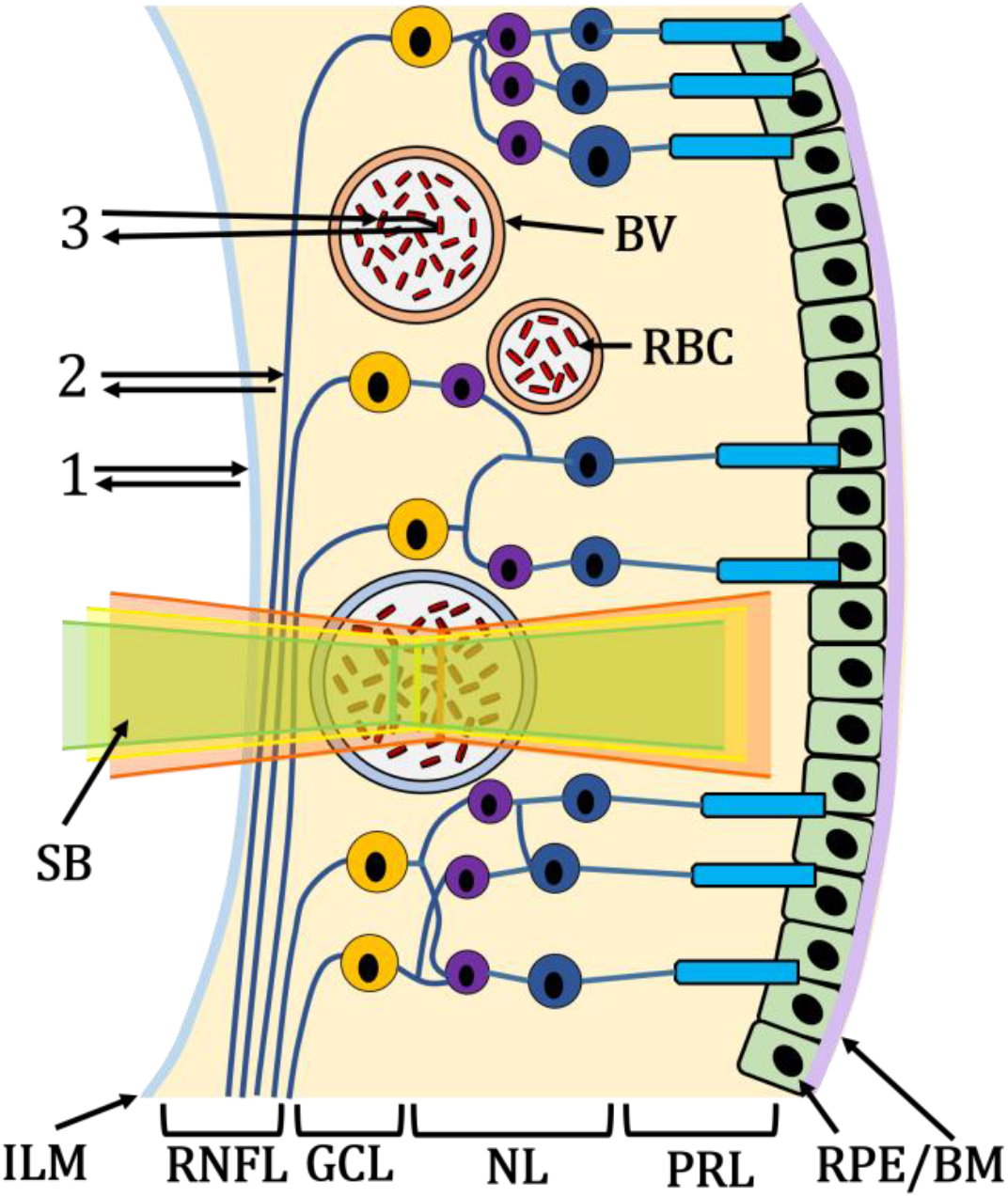
Simplified illustration of the human retina composed of inner-limiting-membrane (ILM), Retinal Nerve Fiber Layer (RNFL), blood vessels (BV; red is the artery, blue is vein), red blood cells (RBC), ganglion cell layer (GCL), nuclear layers (NL) representing the outer nuclear layer to the outer nuclear layer, photoreceptor layers (PRL) containing rods and cones, and the retinal pigment epithelium and Bruch’s membrane (RPE/BM). Number 1 highlights the photon path of a specular reflection, 2 highlights the photon path of backscattering without blood attenuation, 3 highlights the photon path of backscattering with blood attenuation. A scanning beam (SB) is composed of visible-light wavelengths (green, yellow, and red illustrate different spectral bands of the beam).

System-dependent SCs come from the optical illumination, detection, and processing of the vis-OCT signal. We identified three key system-dependent SCs: spectrally-dependent roll-off (SDR), spectrally-dependent background bias (SDBG), and longitudinal chromatic aberration (LCA). Recently, we showed how SDR (27) and SDBG (28) contaminate spectroscopic measurements of *ex vivo* blood samples and *in vivo* humans, respectively, in vis-OCT. LCA (illustrated by the axial displacements of green, yellow, and red wavelengths in the scanning beam in Fig. 1) has also been considered a contaminant for structural and spectroscopic OCT (29-31). Please also refer to **Supplementary Materials – Accounting for LCA**.

Previously, researchers analyzed vis-OCT backscattered signals from a vessel’s posterior wall (PW) to measure sO_2_ in rodents (17, 22) without correcting the SCs mentioned above. The fundamental limitation of a purely backscattering measurement is that it does not directly measure the attenuation coefficient of blood and is therefore susceptible to SCs. This method also was unable to measure sO_2_ from large vessels where the PW was undetectable due to strong blood attenuation.

Both the sample-dependent and system-dependent SCs have magnified and more unpredictable influences on sO_2_ measurement accuracy in human imaging than in small animal imaging because of the reduced light illumination power due to ocular laser safety and patient comfort, stronger eye motion, and larger variation in retinal anatomy (18, 19, 26, 32). In addition, it is more challenging to identify PWs of all human retinal vessels due to enlarged vessel diameters. Therefore, clinical vis-OCT oximetry requires that sO_2_ measurements be free from SCs without identifying PWs of vessels.

In this work, we developed an adaptive spectroscopic vis-OCT, or Ads-vis-OCT, to isolate blood’s spectral signature without needing to identify vessels’ PWs. We validated Ads-vis-OCT’s accuracy in *ex vivo* samples made from bovine blood at 17 oxygenation levels (see **Supplementary – *Ex vivo* phantom verification and comparison**). We tested Ads-vis-OCT’s repeatability in 125 unique vessels from 18 human volunteers imaged in a clinical environment. In our human tests, vis-OCT-measured retinal artery sO_2_ values agreed with the corresponding pulse oximeter readings. Most importantly, our sO_2_ results required no calibration to account for any systemic bias, suggesting an SC-insensitive sO_2_ measurement.

## Results

### Adaptive Spectroscopic vis-OCT

Fig. 2 shows the flow-chart for Ads-vis-OCT, containing 12 steps on how they flow from one to the other. Figures before and after each step depict their respective inputs and outputs. In **Step 1**, we computed the STFT of each full-band interference fringe to obtain 21 spectrally-dependent A-lines (SDA-lines) (see **Methods – Short-time Fourier Transform**). The input is the interference fringes divided into 21 sub-bands, where the colors highlight the center wavelengths (*λ*) in the visible-light spectrum. The outputs are the SDA-lines, which encode depth (z-dimension) and wavelength sub-bands *(λ*-dimension). The SDA-lines experience axial displacement among the 21 sub-bands (highlighted by colored arrows) due to imperfect correction of dispersion mismatch and LCA (33). The SDA-lines also experience a bias from the SDBG (highlighted by black-dashed box).

**Figure 2.**
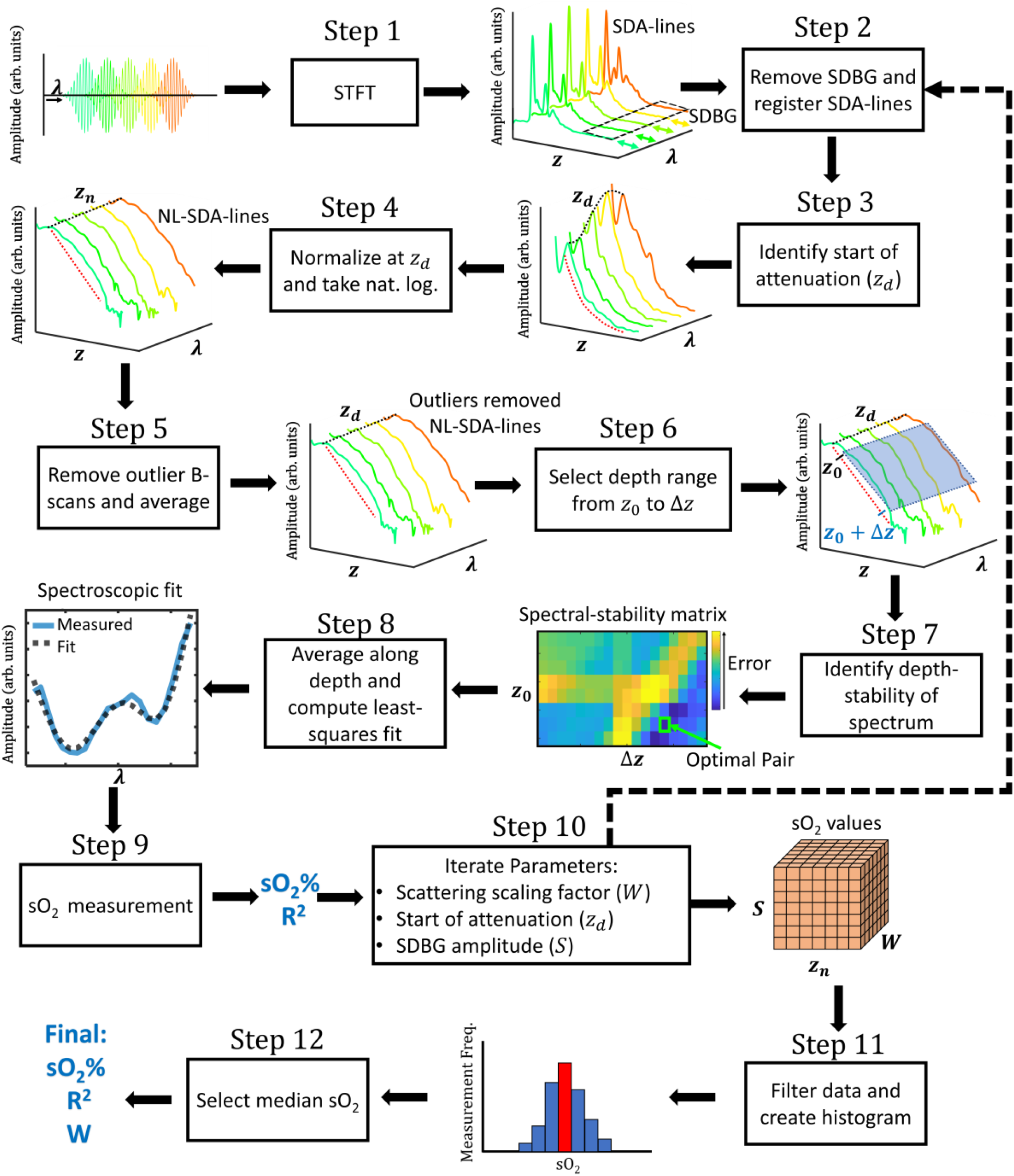
Flow chart overview of Ads-vis-OCT processing for retinal oximetry. Arrow direction highlights the input and output of each step.

In **Step 2**, we axially registered the SDA-lines for all B-scans. We also removed the SDBG bias in the SDA-lines (34). Strategy for registration and SDBG removal are described in **Methods – SDA-line Registration** and **SDBG Removal**.

In **Step 3**, we identified the depth at which signal decay from the blood begins (z_*d*_) in each SDA-line (see **Methods – Normalization Model for Spectroscopic vis-OCT**). As the **Step 3** output, the spectrum at z_*d*_ is highlighted by the black dashed line, and the approximately exponential blood decay is highlighted by the red dashed line.

In **Step 4**, we normalized the SDA-lines by the spectrum at z_*d*_ and computed their natural logarithm, making the original exponential decay almost linear (see **Methods – Normalization Model for Spectroscopic vis-OCT**). In the **Step 4** output, the black dashed line highlights the spectrum at z_*d*_, which is constant across *λ* after normalization. The red-dashed line highlights the approximately linear decay of the vis-OCT blood signal after taking the natural logarithm.

In **Step 5**, we removed outlier B-scans using coarse data filtering (see **Methods – Coarse Data Filtering**). After removing the outliers, we averaged the remaining B-scans to reduce noise, yielding a single set of natural logarithm SDA-lines (NL-SDA-lines).

In **Step 6**, we selected the depths for spectroscopic measurement in the NL-SDA-lines. The **Step 6** outputs are the starting depth (*z*_0_) and depth range (Δ*z*) for the measurement. The blue box highlights the measurement range for selected depths and spectral sub-bands. We averaged NL-SDA-lines in the blue box along the z-axis, yielding a 1D STFT spectrum (see **Methods – Depth Averaging**).

In **Step 7**, we assessed how the STFT spectrum changed along with the depth of the vessel (see **Methods – Depth Selection**). Briefly, we calculated a 1D STFT spectrum at different depths by iterating through different *z*_0_ and Δ*z* values. For each depth, we applied nine perturbations to *z*_0_ and Δ*z*. Then, we calculated changes in the measured spectrum after the perturbations. The **Step 7** output is the spectral-stability matrix (SSM), which plots the shape change (error) after perturbations in *z*_0_ and Δ*z*. We selected the optimal pair of *z*_0_ and Δ*z* with the smallest error (highlighted by green box).

In **Step 8**, we averaged the NL-SDA-lines within the depth range defined by the optimal *z*_0_ and Δ*z* and fit the depth-averaged STFT spectrum to a linear combination of oxygenated and deoxygenated blood attenuation spectra (see **Methods – Oximetry Fitting Model**). The **Step 8** output is the spectroscopic fit, in which a least-squares regression fits a predicted STFT spectrum (black dashed line) to the measured STFT spectrum (blue line).

In **Step 9**, we estimated the sO_2_ value by calculating the proportion of the oxygenated blood in the fit. We also calculated the corresponding coefficient of regression *R*^2^.

In **Step 10**, we adapted to experimental and physiological variables in each vessel by iterating through **Steps 2-9** for small variations in three parameters. These parameters include scattering scaling factor *W*, the start of attenuation depth *z*_*d*_, and amplitude scaling factor *S* in removing SDBG. The outputs of **Step 10** are two three-dimensional (3D) matrixes that store the measured sO_2_ and R^2^, respectively, corresponding to each parameter iteration (see **Methods – Parameter Iterations**).

In **Step 11**, we applied fine data filtering (see **Methods – Fine Data Filtering**), where we created a histogram of the sO_2_ values. In **Step 12**, selected a central value from the histogram. The selected sO_2_ value and its corresponding R^2^ and *W* values were the final outputs.

### Spectroscopic normalization in retinal blood vessels

To extract blood attenuation from vis-OCT signals, we need to remove the sample-dependent SC influences from tissues with depths above the blood as the vis-OCT light propagates (group 1 and group 2 photons highlighted in Fig. 1). Normalization of the blood STFT spectrum by the tissue STFT spectrum at a selected depth can potentially remove sample-dependent SC influences. We tested four normalization methods and compared their corresponding STFT spectra. Fig. 3a is a representative vis-OCT B-scan image acquired 1.7-mm superonasal to the optic disc from a 37-year-old female volunteer. We selected one vein (V1) and one artery (V2) with diameters of 168 µm and 120 µm, respectively. The four normalization methods are Method 1: no normalization, as reported by Yi et al. (17), Chen et al. (25), and Pi et al. (22); Method 2: normalization by the RNFL, which is typically anterior to the retinal vessels, as reported by Song et al. (26) and suggested by Chong et al. (19, 20); Method 3: normalization by the anterior vessel wall (AW), which can be highly reflective and is immediately above the blood signal; and Method 4: normalization by the start of signal decay in the blood (z_*d*_). We applied SDBG correction (28) for all four methods. In Figs. 3b-3i, we plot the measured STFT spectrum for each normalization against the spectrum predicted by least-squares regression (see **Step 8** of Fig. 2). For all methods, we plot the regression’s best fit with respect to normalization method 4, since it is the only one consistent with our theoretical model (see **Methods** – **Normalization Model for Spectroscopic vis-OCT**), experimental verification (see **Supplementary**--***Ex vivo* phantom verification and comparison**), and fitting quality threshold (see **Methods** – **Fine Data Filtering**). If all the normalization methods were equivalent, each measured STFT spectrum would converge to this best-fit spectrum. Below, we show this is not the case.

**Figure 3.**
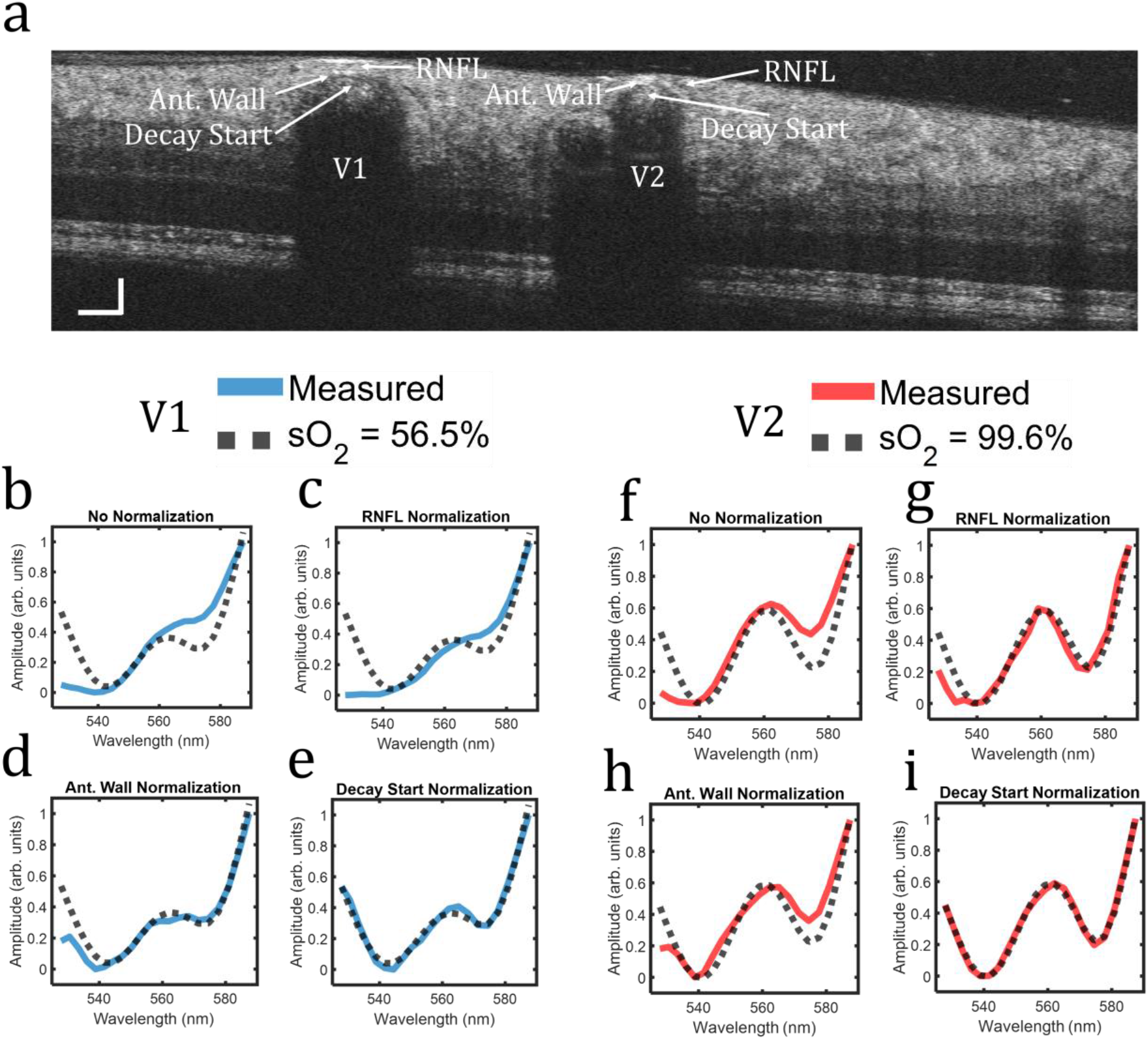
Spectroscopic normalizations in the human retina. (a) vis-OCT B-scan. Vessels labeled V1 and V2 were identified as vein and artery, respectively. Arrows highlight anatomical features used for normalization. (b-e) Measured spectrum (blue line) and best-fit spectrum for sO_2_ = 56.5% (back dashed line) in V1 for no normalization, normalization by the RNFL, normalization by the anterior vessel wall, and normalization by the start of signal decay in blood, respectively. (f-i) The same analysis for the panels b-e is replicated for V2.

In Figs. 3b-3e, we plot the measured STFT spectra (blue lines) in V1 against best-fit spectrum (dashed line) for sO_2_ = 56.5% and scattering scaling factor (17) *W* = 0.05. To standardize the plots, all measured and best-fit spectra are plotted after normalization between 0 and 1. Using Method 1 (Fig. 3b), the measured STFT spectrum overestimates the best-fit spectrum at longer wavelengths and underestimates it at shorter wavelengths. Using Method 2 (Fig. 3c), the measured STFT spectrum follows a similar trend but is not identical to that using Method 1. Using Method 3 (Fig. 3d), amplitudes of the measured STFT spectrum agree well with the best-fit spectrum for wavelengths longer than 540 nm, while they are comparatively lower for wavelengths shorter than 540 nm. Using Method 4 (Fig. 3e), the measured STFT spectrum agrees well with the best-fit spectrum for all wavelengths (R^2^ = 0.99). In addition to removing many of the SC described in Fig. 1, Method 4 is the only one removing influence from the backscattering coefficient of blood. The other three normalization methods each generated measurements that are different from the best-fit result, which means that these methods still suffer from SCs.

Figs. 3f-3i plot spectra measured in V2 alongside the best-fit spectrum for sO_2_ = 99.6% and *W* = 0.09. Using Method 1 (Fig. 3f), the measured spectrum overestimates the best-fit spectrum for longer wavelengths and underestimates it for shorter wavelengths, similar to that in Fig. 3b. In Method 2, since there was no visible RNFL tissue above V2, we normalized by the RNFL tissue directly to the right of the vessel (highlighted by an arrow in Fig. 3a). Using Method 2 (Fig. 3g), the measured spectrum agrees with the best-fit at wavelengths longer than 540 nm but disagrees at wavelengths shorter than 540 nm, similar to the trend seen in Fig. 3d. Using Method 3, (Fig. 3h), the measured spectrum overestimates at longer wavelengths and underestimates at shorter wavelengths. Method 4 (Fig. 3i) generated an STFT spectrum agreeing with the best-fit (R^2^ > 0.99) at all wavelengths, indicating successfully removing SCs.

Fig. 3 suggests that careful normalization is critical for accurately and repeatably extracting the oxygen-dependent attenuation spectrum of blood in the retina. Method 4 is facilitated by direct measurement of blood attenuation from the blood signal. This highlights a critical flaw in identifying PW (17), which indirectly measures blood attenuation from PW backscattering and assumes that it is independent of SCs. We confirmed that normalization Method 4 is the best to remove SCs in other tested vessels, as shown in Fig. S3a in **Supplementary Materials**.

### Accounting for depth-dependent spectra within retinal vessels

The Beer-Lambert model (17, 35) assumes that the spectrum of blood is depth-independent. In reality, we observed changes in the measured STFT spectrum with depth in human retinal vessels, which may be attributed to scattering changes with the vessel depth from the distributions of blood cells (36, 37). Therefore, we developed a spectral stability analysis in **Step 7** (Fig. 1), as detailed in **Methods** – **Depth Selection**. The goal of spectral stability analysis is to select a depth region in a blood vessel where the measured spectrum is mostly depth-independent.

Fig. 4a shows the SSM for V1 from Fig. 3a (see **Methods** – **Depth Selection**). The SSM plots how much the measured STFT spectrum changes with depth as a function of the selected starting depth *z*_0_ and depth range Δ*z*. A lower mean-squared error (MSE) indicates that the measured STFT spectrum has less depth variation. The green box in Fig. 4a highlights the pair of *z*_0_ and Δ*z* where MSE is the lowest, and the black box highlights where it is highest. We plot the measured STFT spectra corresponding to the black and green boxes in Fig. 4a in **Supplementary Materials** (Fig. S7a). In V1, the spectrum is most depth-stable for longest Δ*z* and least depth-stable for shortest Δ*z*. Optimal depths varied across the veins investigated in this study. The average selected Δ*z* for veins (n = 53) was 33 *μ*m out of a maximum value of 40 *μ*m. Selected Δz ranged from Δz = 22 *μ*m to Δz = 40 *μ*m.

**Figure 4.**
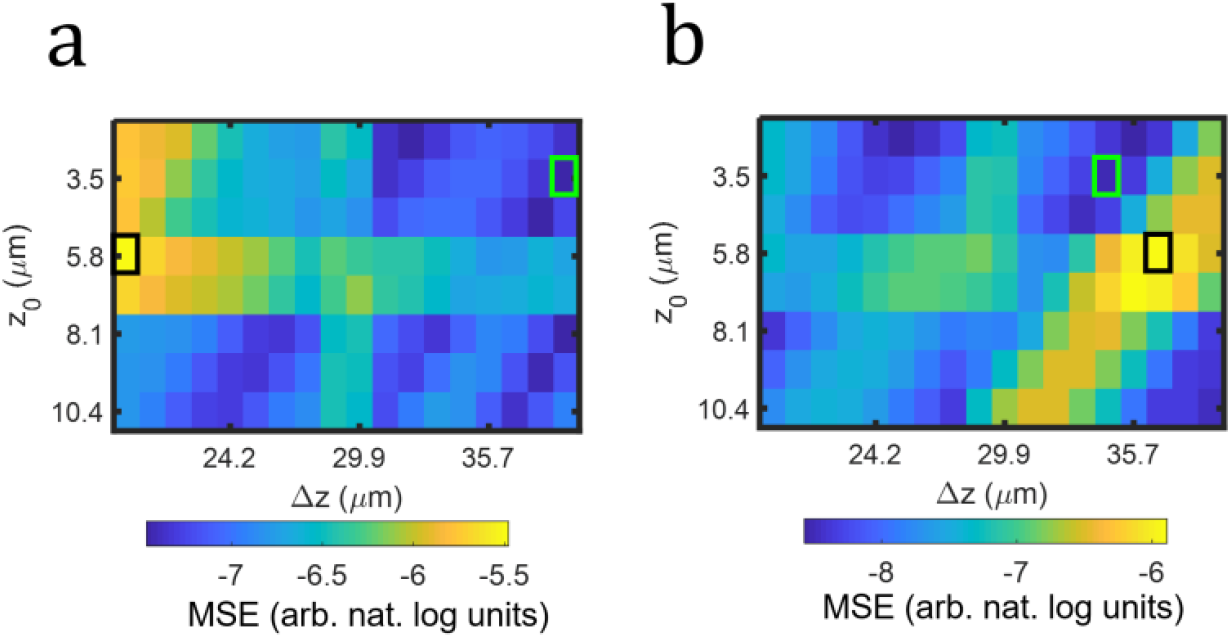
SSM for (a) V1 and (b) V2 in Fig. 3. The green box highlights the lowest MSE and the black box highlights the highest MSE.

Fig. 4c shows the SSM for V2 from Fig. 3a. Unlike Fig. 4a, the lowest MSE (green box) is actually at a shorter depth than the highest MSE (black box). Arteries are generally pulsatile and have higher flow velocity than veins, perhaps introducing different SC with the vessel depth. Nevertheless, we show that by selecting the *z*_0_ and Δ*z* with the lowest MSE, we selected a depth-stable spectrum. We plot the measured STFT spectra corresponding to the black and green boxes in Fig. 4b in **Supplementary Materials** (Fig. S7b). Similar to veins, the most and the least stable depth varied across arteries in this study. The average selected Δ*z* in arteries (n = 72) was 33 *μ*m out of a maximum of 40 *μ*m and selected Δ*z* ranged from Δ*z* = 17 *μ*m to Δ*z* = 40 *μ*m. We analyzed the spectral stability for other selected vessels in the **Supplementary Materials**.

### Retinal oximetry around the optic disk

The network of vessels supporting oxygen delivery to the inner retina is derived from the optic disk. Therefore, an oximetry map of the optic disk can help to investigate oxygen delivery or extraction in the entire retina. We achieved an oximetry map of the optic disk with a single 10-s vis-OCT scan. Fig. 5a illustrates a representative oximetry map of the optic disk with a 4.8×4.8 mm^2^ FOV in a 23-year-old volunteer without known ocular diseases. We measured sO_2_ in 17 vessels (10 arteries and 7 veins) ranging from 37 µm to 168 µm in diameter.

**Figure 5.**
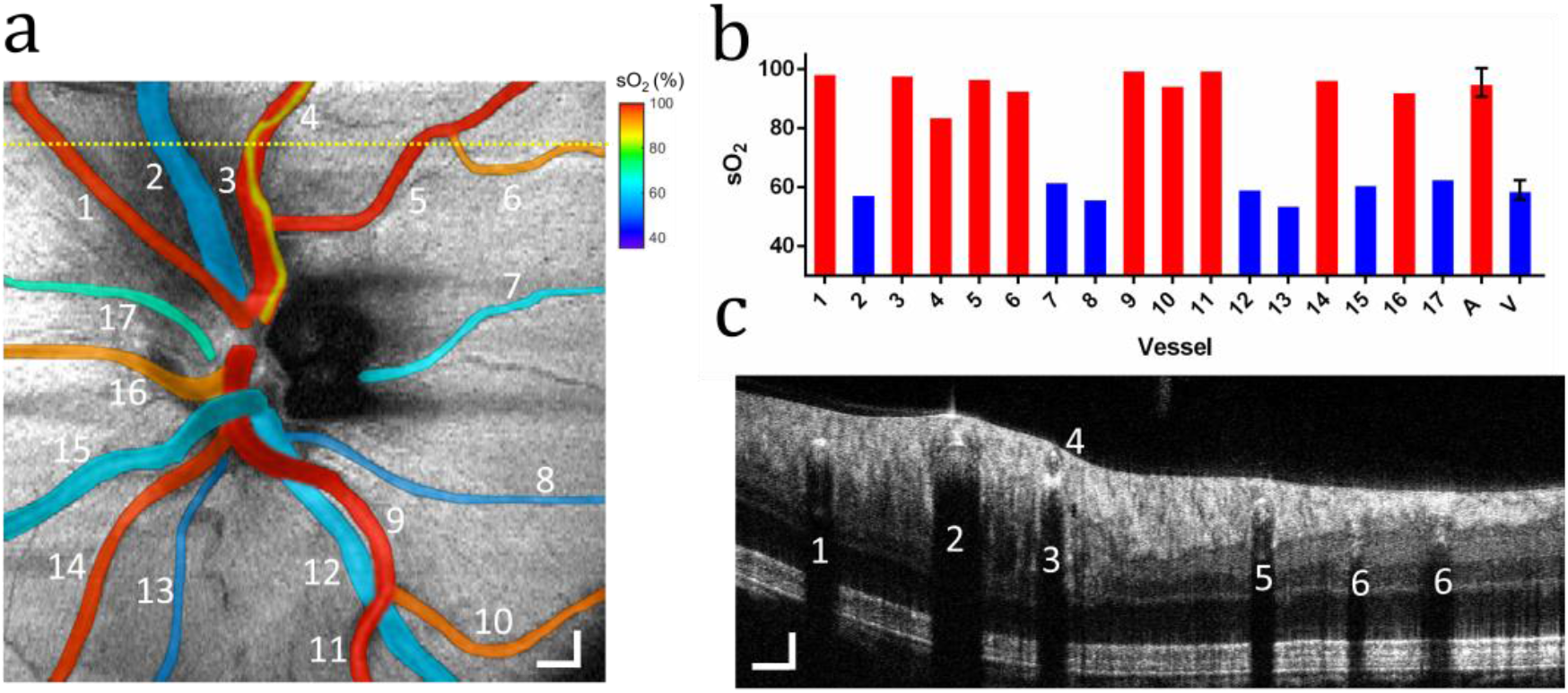
Oximetry map of the optic disk. (a) sO_2_ measurements in 17 vessels in the optic disk from a healthy 23-year-old volunteer. The sO_2_ values pseudo-colored and overlaid onto the fundus image. Scale bar: 300 *μ*m; (b) Bar chart plots sO_2_ measurements from the panel a in individual arteries (red bar) and veins (blue) numbered from 1 to 17, as well as average sO_2_ in all arteries (A) and all veins (V); (c) B-scan from the position highlight by the yellow dashed line in panel a. Scale bars: 150 *μ*m.

We pseudo-colored the 17 vessels according to their measured sO_2_ values onto a vis-OCT fundus image (Fig. 5a) and plotted the sO_2_ values in the bar chart (Fig. 5b). The measured sO_2_ across all arteries was 95.8±4.4% (n = 10) and the measured sO_2_ across major arteries (diameter ≥ 100 µm) was 97.3%±2.8% (n = 6). The average pulse oximeter measurement from the index finger was 98%, which agrees well with the vis-OCT measured sO_2_ from the major arteries. The measured sO_2_ across all veins was 59.0±3.2%, and the mean sO_2_ between major arteries and veins (A-V difference) was 38.3%.

Fig. 5c shows a B-scan from the location highlighted by the dashed yellow line in Fig. 5a. We can observe a small artery (vessel 4, diameter = 37 µm) directly above a major artery (vessel 3, diameter = 122 µm). The measured sO_2_ value in vessel 3 is 98.3%, consistent with pulse oximeter reading (98%), and the measured sO_2_ value in vessel 4 is 85.8%. We were able to measure sO_2_ values from both vessels (Fig. 5a), demonstrating the unique depth-resolved sO_2_ imaging capability permitted by vis-OCT. Such axially overlapping vessel anatomy further emphasizes the need for AdS-vis-OCT since the oxygen-dependent spectrum from vessel 4 will contaminate the measured blood spectrum from vessel 3 if not correctly normalized. Our AdS-vis-OCT measured sO_2_ in both vessels, independent of one another. The posterior wall of vessel 4 appears to be in direct contact with the anterior wall of vessel 3. We hypothesize that the lower measured sO_2_ in vessel 4 is partially caused by oxygen diffusion to the contacting vessel or surrounding RNFL.

### Retinal oximetry in a healthy cohort

We performed vis-OCT retinal oximetry in 18 volunteers without known health issues in clinics. We measured sO_2_ in 125 unique vessels (72 arteries and 53 veins) within a 3.4 mm radius of the optic nerve head.

Fig. 6a shows sO_2_ from unique arteries (red) and veins (blue) plotted as a function of vessel diameter. Arterial sO_2_ values show a decreasing trend with decreasing vessel diameter. We determined that vessels diameter was a statistically significant factor (see **Methods – Statistical Analysis**) in this trend (*p* = 4.35 × 10^−6^). The diameter-dependent trend is consistent with oxygen gradients observed in other precapillary arteries (38-44). Since smaller vessels generally offered fewer pixels in vis-OCT images to average and therefore were potentially more sensitive to noise, we investigated whether the sO_2_ decrease was an artifact of lower spectral fit R^2^. We determined that R^2^ was not a statistically significant factor (see **Methods – Statistical Analysis**) in this trend (*p* = 0.701). Venous sO_2_ slightly increases with decreasing vessel diameter, but the trend is not statistically significant (*p* = 0.232). Spectral fit R^2^ is also not significant (*p* = 0.070) in determining sO_2_ in veins.

To account for the observed sO_2_ gradient with diameter, we computed average sO_2_ in arteries across two diameter groups (diameter ≥ 100 µm and diameter < 100 µm). Fig. 6b shows sO_2_ measurements for major arteries (diameter ≥ 100 µm), small arteries (diameter < 100 µm), and veins with all diameters. Major arteries (n = 36) had sO_2_ = 97.9 ± 2.9%. Small arteries (n = 36) had sO_2_ = 93.2 ± 5.0%. The difference in sO_2_ between the two groups was statistically significant (*p* = 4.01 × 10^−6^, two-sample T-test). Average spectral fits were R^2^ = 0.96, 0.93, and 0.95 for major arteries, small arteries, and all veins, respectively. An average R^2^ of 0.93 or higher validates the efficacy of Ads-vis-OCT in a wide range of retinal vessel diameters in humans.

**Figure 6.**
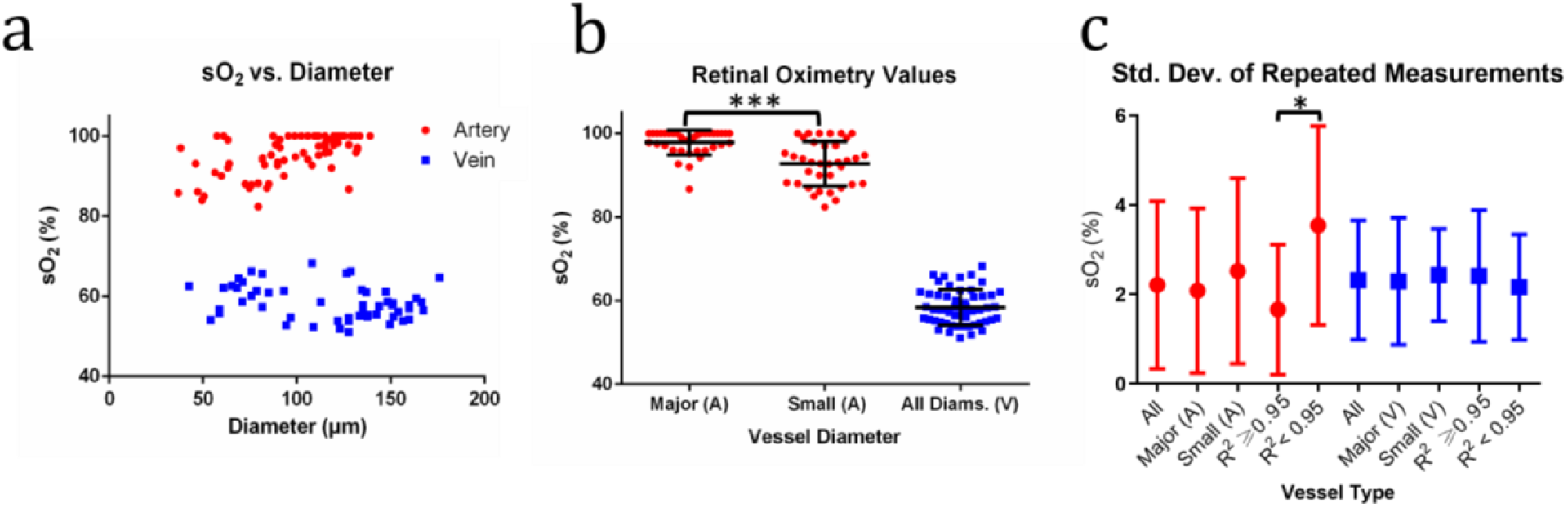
Retinal sO_2_ in a healthy cohort. (a) sO_2_ measurements in 72 unique arteries (red) and 53 unique veins (blue) plotted against vessel diameter from 18 healthy volunteers; (b) Distribution of sO_2_ measurements for major (diameter ≥ 100 *μm*) and small (diameter < 100 *μm*) artery calibers and all veins; (c) Repeatability of arteries and veins. * indicates *p* < 0.05 and *** indicates *p* < 0.001 from two-sample t-test.

We acquired repeated scans of 42 unique vessels and calculated their average SDs (24 unique arteries and 18 unique veins across 12 volunteers). Fig. 6c shows average SDs for arteries (red) and veins (blue). All arteries and veins had average SDs of 2.21% and 2.32%, respectively. We noted above that smaller vessels endured less volumetric averaging than larger vessels. Therefore, we investigated repeatability for arteries and veins of diameters larger and smaller than 100 *μ*m. Larger arteries (n = 17) and smaller arteries (n = 7) had average SDs of 2.08% and 2.52%, respectively. Larger veins (n = 15) and smaller veins (n = 3) had average SDs of 2.29% and 2.43% respectively. There was no statistically significant difference between the average repeatability values for any of the groups (two-sample T-test). Finally, we investigated repeatability for best spectral fits (R^2^ ≥ 0.95) and relatively lower spectral fits (R^2^ < 0.95). Best fit arteries (n = 17) and lower fit arteries (n = 7) had average SDs of 1.66% and 3.54%, respectively. Best fit veins (n = 11) and lower fit veins (n = 7) had average standard deviations of 2.41% and 2.16%, respectively. The difference between the average SDs for the artery groups was statistically significant (*p* = 0.02). However, one analyzed artery with R^2^ < 0.95 was an outlier (SD = 6.83%). After removing the outlier, the average SD for R^2^ < 0.95 was 2.99%, and the difference between the best fit and lower fit groups would no longer be statistically significant (*p* = 0.09). Cumulative results for arteries and veins suggest high repeatability in the data acquired in the clinical setting. Statistics for each unique vessel in repeatability analysis shown in Table S1 (see **Supplementary Materials**).

We estimated the ground-truth retinal sO_2_ in major arteries (diameter ≥ 100 um) using the sO_2_ measured from a pulse oximeter. Table 1 compares measured sO_2_ from the pulse oximeter and vis-OCT for unique major arteries (diameter ≥ 100 um) from 12 volunteers, where a pulse oximeter concurrently measured sO_2_ at their index fingers. We measured the root-mean-squared-error (RMSE) between pulse oximeter sO_2_ and vis-OCT sO_2_ for each unique artery in each respective subject (Table 1, Column 5). The last row in Table 1 shows average results across all 12 subjects. Average major artery was sO_2_ = 98.3%, in close agreement with that by the pulse oximeter (sO_2_ = 98.6%). The average RMSE between vis-OCT and the pulse oximeter was 2.04%. Such accuracy is consistent with *ex vivo* vis-OCT sO_2_ measurements (Fig. S2a in **Supplementary Materials**). We also noted that the SD for major arteries was 2.08% (Fig. 6c), which matches the average RMSE. This observation suggests that vis-OCT measured sO_2_ may be within noise-limited agreement with the pulse oximeter. We emphasize that vis-OCT sO_2_ measurements agreed with the pulse oximeter without any post-hoc calibrations (25, 45).

**Table 1.** Vis-OCT retinal oximetry comparison with the pulse oximeter readings. RMSE indicates root-mean-squared error between unique major arteries in each subject and respective pulse oximeter readings.

| Subj. | Avg. Pulse Ox. sO<sub>2</sub> (%) | Avg. vis-OCT sO<sub>2</sub> (%) | Num. Unique Major Arteries Measured by vis-OCT | RMSE Between Pulse Ox. and Unique Artery sO<sub>2</sub> (%) |
| --- | --- | --- | --- | --- |
| 1 | 100 | 100 | 2 | 0 |
| 2 | 98 | 99.6 | 2 | 1.61 |
| 3 | 99.7 | 98.9 | 3 | 1.99 |
| 4 | 99 | 95.2 | 1 | 3.8 |
| 5 | 95 | 95.8 | 1 | 0.8 |
| 6 | 98 | 100 | 1 | 2 |
| 7 | 99 | 100 | 1 | 1 |
| 8 | 100 | 98 | 2 | 2.83 |
| 9 | 98 | 100 | 2 | 2 |
| 10 | 100 | 96 | 2 | 4 |
| 14 | 98 | 97.6 | 5 | 2.59 |
| 17 | 99 | 99 | 3 | 1.38 |
| Avg. | 98.6 | 98.3 | 2.1 | 2.04 |

### Comparison of depth-averaging and slope methods

The depth-resolved slope of NL-SDA-lines (further referred to as the “slope method”) was previously used to extract the attenuation coefficient of OCT signals in **Step 8** in Fig. 2 (19, 29, 35). In this work, we found that the depth-resolved average of NL-SDA-lines (further referred to as “depth-averaging method”) was superior to the slope method for retinal oximetry. We compared the two methods for sO_2_ measurements in the 125 unique human retinal vessels described above. Both sO_2_ measurements used identical AdS-vis-OCT processing with identical depth-selection windows. To implement the slope method, only the depth-averaging step, as depicted by Step 8 in the AS-OCT processing, was replaced with a simple linear regression to estimate the slopes of NL-SDA-lines (output of Step 6) along the *z*-axis.

Table 2 compares the sO_2_ measurement statistics for 125 unique human retinal vessels (72 arteries and 53 veins) for the depth-averaging and slope methods. Success rate indicates the fraction of sO_2_ measurements that surpassed our quality control threshold, including a spectral fit R^2^ > 0.80 or R^2^ > 0.93 if sO_2_ = 100%. For this analysis, we combine sO_2_ across all diameters.

**Table 2.**
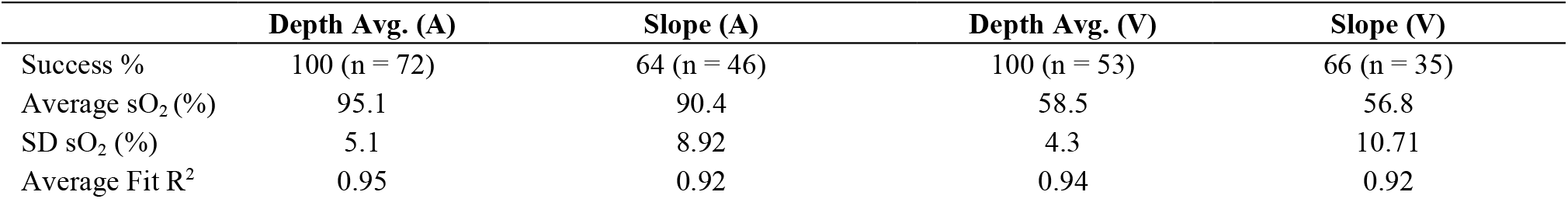
Comparison of depth-averaging and slope methods for vis-OCT retinal oximetry. (A) indicates artery and (V) indicates vein.

For all unique arteries, the depth-averaging method had a success rate of 100 %, while the slope method had a success rate of 64 %. For depth-averaging, all arteries had an average sO_2_ of 95.1 ± 5.1 %, while the slope method yielded an average of 90.4 ± 8.92 %. The SD of the slope method was 75% higher than that of the averaging method. The lower average spectral fit R^2^ for the slope method (0.92), as compared with the depth-averaged method (0.95), is consistent with its higher SD. Additionally, we anticipated that the R^2^ was still inflated for the slope method since we rejected 36% of vessels based on a low R^2^.

For all unique veins, the depth-averaging method had a success rate of 100%, while the slope method had a success rate of 66%. For depth-averaging, veins had an average sO_2_ of 58.5 ± 4.3 %, while the slope method had an average of 56.8 ± 10.71 %. SD of the slope method was 149% higher than that of the depth-averaging method. Similar to arteries, the slope method had a lower average R^2^ (0.92), as compared with the depth-averaging method (0.94). Since we rejected 34 % based on low R^2^, we anticipate that the venous R^2^ for the slope method was still inflated.

The comparisons between the two methods demonstrate that the depth-averaging method greatly improves the stability of vis-OCT retinal oximetry than the slope method. We investigated the improved stability using Monte-Carlo simulation in the **Supplementary Materials**.

## Discussion

Clinical application is the ultimate goal for vis-OCT oximetry, which SC currently hinders. In this work, we developed depth-resolved Ads-vis-OCT processing to remove SCs from sO_2_ measurements. We also introduced a depth-averaging method to measure attenuation spectra that significantly improved stability for sO_2_ measurements *in vivo*.

Using Ads-vis-OCT, we found excellent agreement with reported spectra *ex vivo* and *in vivo* (Fig. S2). In the human retina, sO_2_ measurements across all unique arteries and veins were highly repeatable (average SD = 2.21% and 2.31%, respectively). Furthermore, the average error between major artery sO_2_ and a pulse oximeter (RMSE = 2.08 %) was similar to the average SD of major artery sO_2_ after repeated measurements (SD = 2.04 %), suggesting that the accuracy was limited by noise and not a systemic bias. Across 72 unique arteries from 18 subjects, we found a statistically significant trend between decreased diameter and decreased sO_2_. This trend is consistent with previously observed precapillary oxygen gradients (38-44).

Previous vis-OCT retinal oximetry relied on backscattered signal from the vessel PW to measure sO_2_. The PW method is an indirect measurement of blood’s attenuation spectrum and does not consider SCs other than the PW itself. Contamination by the PW is fit to a power-law decay (*Bλ*^-*α*^) by the first Born approximation (17), where *B* and *α* are positive constants representing microsctructural scattering properties of the tissue. As previously recognized (45), and illustrated by the measured STFT spectra in Fig. 3, such an sO_2_ model is likely overly simplified, especially in the human retina. First, most vessels are buried under several layers of retinal tissues with varying optical properties (14, 19, 32, 46, 47). Interfaces at the ILM and vessel wall are composed of highly reflective and fibrous tissues that may not necessarily conform to power-law approximation. For example, consider vessel V1 in Fig. 3. The ILM/RNFL interface appears transparent at some parts of the retina but is bright directly above the blood vessel. Since the brightness is most intense when the interface is orthogonal to the vis-OCT illumination beam, we hypothesize that it may come from specular reflection or high backscattering from the fibrous tissues. A similar trend is valid at the anterior wall (AW) interface for V1. While the majority of the vessel wall has similar intensity to the RNFL, the center of the AW exhibits a higher reflectance. We believe this is either specular reflection or perhaps backscattering from the fibrous vessel lamina (47). Such light-tissue interactions can have spectral profiles dependent on the incident angle of light, as well as local optical properties such as polarization (48, 49). As another example, the ILM in Fig. S5 (see **Supplementary Materials**) shows higher amplitudes for shorter wavelengths (green) than longer wavelengths (orange), but the reverse trend at the vessel wall, which is not consistent with the power-law decay model. In general, the non-blood tissues examined in our study did not exhibit reliably identical structural or spectral features to apply such a simplified decay model. We hypothesize previous uses of the power-law model may have partially fit other SCs.

Song et. al previously attempted to remove some SCs using the PW method by normalizing the measured STFT spectrum by signal backscattered from the RNFL (26). As seen in Eqns. 1-7, normalizing by tissues other than the blood itself results in contaminating terms that do not isolate the attenuation spectrum of blood. This is supported by Fig. 3, where we show that normalizing by different tissues results in different measured spectra. By directly measuring blood attenuation and normalizing at *z*_*d*_, we show analytically (Eqns. 1-7), *ex vivo* (**Supplementary** Fig. S2), and *in vivo* (Fig. 3) that most SCs can be removed, thereby providing consistent and accurate measurements. Direct blood measurement also enables sO_2_ measurements in large and small vessels independent of visibility of the PW.

Vis-OCT measures a scattering coefficient of blood lower than that predicted by Mie theory (23). Here, we represent its reduction by multiplication with the coefficient *W*. However, the value of *W* in vis-OCT and the reasoning for its existence are still not well agreed upon and varies greatly depending on the studies (17, 20, 29, 50). Several vis-OCT works used *W* = 0.2, which is based on a model dependent on hematocrit (17). To our knowledge, this value was derived *in vivo* in rat retinas without removal of SCs we identified in this work. Following the appropriate normalizations in a well-controlled *ex vivo* phantom experiment (**Supplementary** Fig. S2), we found strong fits between *W* = 0.02 and *W* = 0.10, nearly 10-fold smaller than the reported *W* = 0.2. The average best-fit *W* in the phantom experiment was 0.068 at physiological hematocrit (45%) (Fig. S2a). Since our normalization protocol explicitly isolated the scattering and absorption coefficients of blood, we anticipated that spectral measurements in the human retina should be highly similar to those *ex vivo* (neglecting any spectral differences between human and bovine blood). We measured the average best-fit *W* in the human retina as 0.064 (Fig. S2c), nearly identical to that found in the *ex vivo* experiment (0.068). Even though the two experimental conditions and sample media were very different, we reached nearly identical quantitative conclusions. We believe that the prior conclusion of *W* = 0.2 was the result of incomplete normalization of the blood spectrum since a higher *W* value results in spectral shapes similar to those shown without normalization in Fig. 3 (17). To this end, the higher *W* may potentially fit SCs from the vis-OCT system or tissue, but not the true attenuation coefficient of blood. Misinterpreting SCs for blood attenuation can result in overfitting of oxygen-dependent parameters, inflating spectral fit R^2^, and leading to overconfident or inaccurate measurements.

Recent vis-OCT measurements of oxygenated hemoglobin by Veenstra et al. (29) recognized many of the system-dependent spectra depicted here. They found that the average scattering coefficient of blood was significantly reduced to ∼100 cm^-1^, equivalent to *W* = 0.03 and within our observed *W* range. We note that they did not consider oxygen-dependent scattering by the Kramers-Kronig relationship (23), which our work does. As Veenstra et al. noted, the reduced scattering contribution is likely caused by the high forward cattering of erythrocytes. Such forward scattering increases the likelihood that multiply scattered photons are collected within the illumination beam’s solid angle. Since the multiply scattered photons will reach deeper than singly scattered photons, the perceived scattering coefficient in the blood is reduced.

Vis-OCT signal amplitudes in the blood are scaled by the backscattering spectra of erythrocytes (51) (Eqs. 1&2 in **Methods**). A previous study developed a theoretical model for erythrocyte backscattering (52) in the visible-light spectral range, although it did not find strong agreement with experimental measurements. In this work, we directly measured and normalized the influence of blood’s backscattering spectrum, which is distinct from its attenuation spectrum.

Additionally, the reduced axial resolution of spectroscopic vis-OCT analysis may prevent delineation of the backscattered blood signal from other tissues in small vessels and capillaries. Recent work by Pi *et al*. (22) reported an STFT central wavelength of 555 nm with a bandwidth of 9 nm. Assuming a refractive index of 1.35, this provides an *in vivo* STFT axial resolution of ∼11 µm. Although this work segmented vessels using the full-band axial resolution of vis-OCT (reported 1.2 µm), their spectroscopic measurements were limited to the STFT axial resolution. It would be challenging to differentiate contaminating tissue or blood backscattering signals from blood attenuation signals, considering that capillary diameters are comparable to or even smaller than the STFT axial resolution. However, their model for capillary-scale sO_2_ measurement was based on the previously published PW method, which, in addition to not addressing the resolution problem, did not consider SCs for accurate measurement of optical properties of blood. When vessels become sufficiently small (e.g., diameters < 20 µm) such that there is insufficient depth or depth resolution to isolate attenuation contrast, the blood backscattering spectra, rather than attenuation spectra, may be key for accurate sO_2_ measurements, as previously suggested by Liu *et al*. (50).

In summary, we leveraged the high-resolution, depth-resolved advantages of vis-OCT towards the isolation of spectral signatures from light-blood interactions. We developed and tested Ads-vis-OCT for retinal oximetry in 18 healthy subjects in vessels ranging from 37 µm to 176 µm in diameter in a clinical environment. We found excellent spectral fits, repeatability, and agreements with the pulse oximeter readings. AdS-vis-OCT sets the stage for clinical vis-OCT retinal oximetry.

## Methods

### Short-time Fourier transform

To extract a depth-resolved spectrum, we multiplied 21 Gaussian windows with the spectral interferogram. Windows were of equal wavenumber (*k*) full-width at half maximum (FWHM) and spaced equidistantly in the *k* space from 528 to 588 nm. Window FWHM was 11 nm at 558 nm, reducing the axial resolution to ∼ 9 μm in the retina (assuming a refractive index of 1.35). We computed the STFT for the Gaussian windows to generate SDA-lines.

### Normalization model for spectroscopic vis-OCT

To accurately quantify sO_2_, the oxygen-dependent attenuation spectrum of blood must be correctly isolated from sample-dependent and system-dependent SCs. We developed a model of the SDA-line to include alterations induced by the SC on the attenuation spectrum of blood. Eq. 1 describes the SDA-line for a homogenous medium

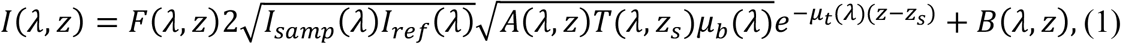

where *λ* is the wavelength; *z* is the depth coordinate; *z*_*s*_ is the surface of the medium with respect to the zero-delay depth 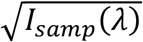 and 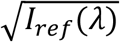 are the power spectra of the light collected from the sample and reference arms, respectively. System-dependent SC spectra include the SDR *F*(*λ, z*) (27), the LCA transfer function *A*(*λ, z*), and the SDBG *B*(*λ, z*) Sample-related SC spectra include the backscattering coefficient of the medium *μ*_*b*_(*λ*), the attenuation coefficient of the medium *μ*_*t*_(*λ*), and he double-pass transmission coefficient across the top interface of the medium *T*(*λ, z*_*s*_).

The retina-specific model for the SDA-line must consider multi-layered media with different optical properties. After normalizing the source power spectrum and subtracting the SDBG, we write the SDA-line at the boundary (*z*_*d*_), where signal decay begins as

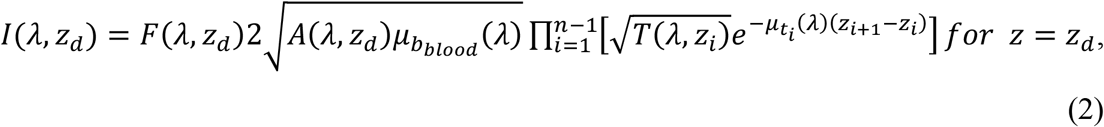

where *i* is the tissue layer and blood is the *n*^*th*^ tissue layer. Furthermore, we write the residual SDA-line below *z*_*d*_ as

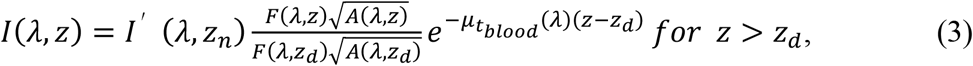

where 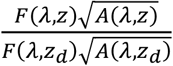 represents the residual LCA and SDR beyond the depth *z*_*d*_. We determined *z*_*d*_ by automatically finding the maximum signal intensity and the starting depth of decay inside the vessel lumen. SDA-lines (Fig. S5) from a vein are representative of the multilayered features described by Eqs. 2 and 3. To reduce noise variation from speckle and background, we calculated *I*′(*λ, z*_*d*_) by depth-averaging the SDA-line across a 6-*μm* region centered at *z*_*d*_. We normalized *I*(*λ, z*) by *I*′(*λ, z*_*d*_) to yield

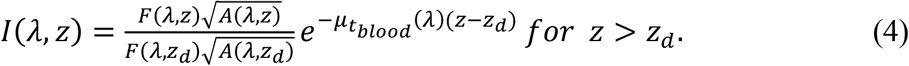

We rejected all vessels from depths greater than 800 µm from the zero-delay. We calculated 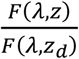 from the roll-offs of our spectrometer up to 65 *μ*m into the vessel and found the SDR had negligible spectral influence after normalization by *I*′(*λ, z*_*d*_). Therefore, the ratio 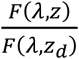 is set to 1, yielding

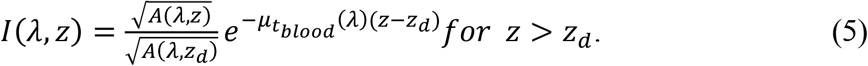

We estimated 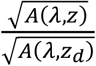 for the same depths (see **Supplementary Materials – Accounting for LCA**) and concluded that the residual LCA could have a small, but a non-negligible influence on sO_2_ even after normalization by *I*′(*λ, z*_*d*_). We, therefore, included 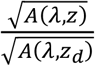 our model. Finally, taking the natural logarithm of *I*(*λ, z*), we have a function that is linearly proportional to 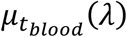 (Fig. 2, Step 3) as

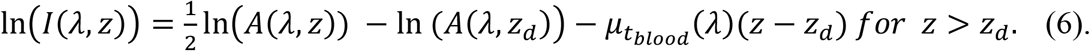

### Coarse data filtering

We averaged NL-SDA-lines along a 32 *μ*m depth region beyond *z*_*n*_ for each respective B-scan to obtain a 1-dimensional (1D) STFT spectrum. We calculated sO_2_ and spectral fit R^2^ from NL-SDA-lines in each B-scan by least-squares fit (see **Methods – Oximetry fitting model**). Then, we applied a threshold of sO_2_ > 15% and R^2^ > 0.40 for 1D spectra extracted from each B-scan and removed B-scans that did not pass. We averaged the NL-SDA-lines across all passing B-scans to further reduce noise. For the smallest analyzed vessels (diameter ≤ 60 *μ*m), we did not perform coarse data filtering, since averaging fewer pixels often made the spectroscopic signals too noisy for analysis in individual B-scans. Instead, we directly averaged these B-scans.

### Depth averaging

To remove noise variations in the speckle and the background, we averaged Eq. 6 across *z*, which can be written as

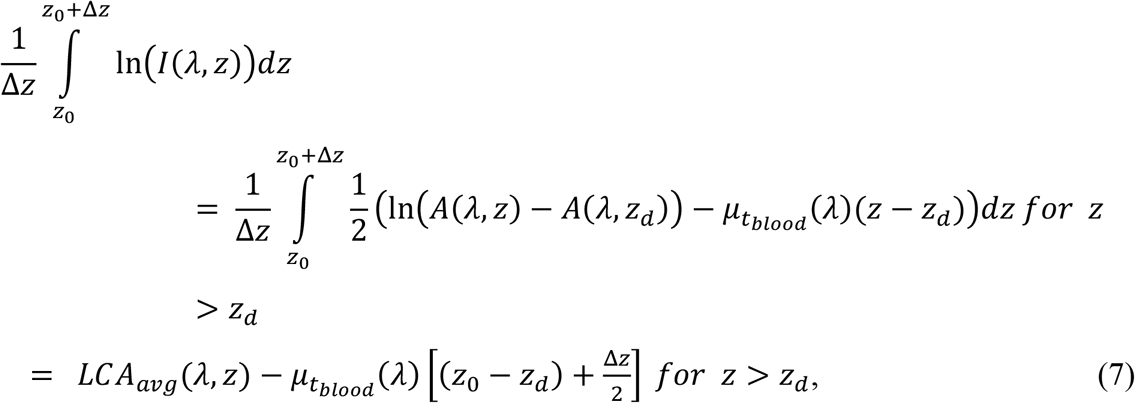

where *LCA*_*avg*_ is the depth-averaged residual LCA; *z*_0_ is the starting depth for averaging; and Δ*z* is the depth range for averaging. The result of Eq. 7 is a linear combination of the *LCA*_*avg*_ and 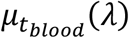. *LCA*_*avg*_ is depicted in Fig. S4 in **Supplementary Materials**.

### Depth selection

Assuming 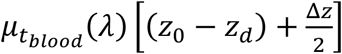 is the dominant form of spectral contrast in Eq. 7 (see **Supplementary Materials – Accounting for LCA**), in the ideal case, the spectral shape of the measured STFT spectrum should not change with depth. However, we observed that the measured STFT spectral shape occasionally changed with depth. To minimize these effects, we elected to analyze *z*_0_ and Δ*z* where the measured STFT spectrum changed the least in response to perturbations in depth. To assess this condition, we developed the spectral stability matrix (SSM).

First, we measured the STFT spectrum in a blood vessel according to Eq. 7. We iterated *z*_0_ from 0 to 12 *μm* and Δ*z* from 17 to 40 *μm*, both in 1.15 *μm* (depth pixel size) increments. We normalized each spectrum to a minimum of 0 and a maximum of 1. Then, we generated a 3-dimensional (3D) matrix that indexed the spectra according to their respective depth windows. Such a matrix can be written as

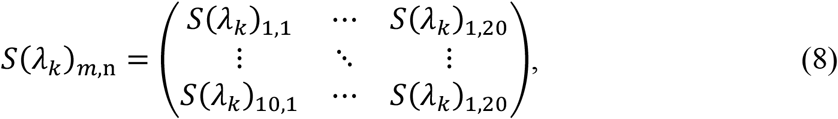

where *S*(*λ*_*k*_)_*m,n*_ is the normalized (between 0 and 1) spectrum for each window iteration; *m* is the iteration index of *z*_0_; *n* is the iteration index of Δ*z*; and *k* is the STFT sub-band index. To measure the response of *S*(*λ*_*k*_)_*m,n*_ to a depth perturbation, we computed the mean-squared-error (MSE) between spectra from 9 adjacent windows in *S*(*λ*_*k*_)_*m,n*_ to generate the SSM as

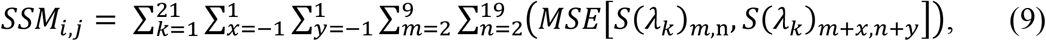

where *MSE*[*S*(*λ*_*k*_)_*m,n*_, *S*(*λ*_*k*_)_*m*+*x,n*+*y*_] is the MSE between spectra *S*(*λ*_*k*_)_*m,n*_ and *S*(*λ*_*k*_)_*m*+*x,n*+*y*_; and *x* and *y* are the indexes of the compared spectra. We show an example *SSM*_*m,n*_ from two selected vessels in Fig. 4. We identified the indexes *m* and *n* where *SSM*_*m,n*_ was minimal the corresponding *z*_0_ and Δ*z*. We used this depth window (e.g., green boxes in Figs. 4a and 4b) for sO_2_ calculation.

### Oximetry fitting model

To extract sO_2_, we fit the spectrum determined by Eq. 7 and the SSM to the following model using a non-negative linear least-squares regression

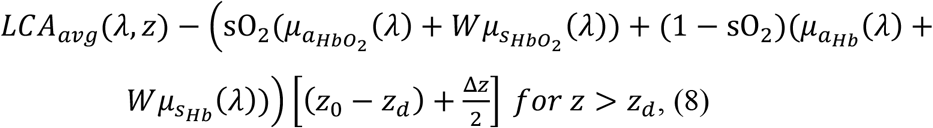

where 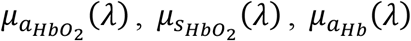, and 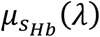 are the reported absorption and scattering coefficients of oxygenated and deoxygenated blood, respectively. *W* is the scattering scaling factor, which scales the above scattering coefficients (17). We computed the fitting for 0.02 ≤ *W* ≤ 0.10 (see **Supplementary Materials — *Ex vivo* phantom verification and comparison**).

The contribution of *LCA*_*avg*_(*λ, z*) in Eq. 8 was estimated using optical simulation and fit the measured spectrum (see **Supplementary Materials – Accounting for LCA**). We accepted the *LCA*_*avg*_(*λ, z*) and sO_2_ value that generated the highest spectral *R*^2^ fit.

We used the reported absorption and Mie-theory-predicted scattering coefficients of blood (14, 23). We modified the reported spectra to match the post-processing of the vis-OCT signal. First, we cropped wavelengths from the reported spectra within our spectrometer’s wavelength range (508 nm – 614 nm). We upsampled the reported spectra by 6-fold to a 12288 element array, which is the same size as our interference fringe after 6-fold zero-padding. Then, we performed interpolation of the reported spectra to be linear in *k*-space, matching the interference fringe’s interpolation. Finally, we filtered and digitized the reported blood spectra with the same 21 STFT Gaussian windows.

### Parameter iterations

We recognized that sO_2_ measurements might be susceptible to minute experimental or physiological variations. We perturbed sO_2_ measurements by introducing minor variations in three parameters we identified as sensitive to sO_2_ accuracy. The first parameter is z_*d*_, the identified depth where blood signal begins to decay. After STFT, the axial resolution of ∼ 9 *μm* and spatial averaging between B-scans broadens the peak blood backscattering signal, adding uncertainty to its localization. Furthermore, random or unknown parameters, such as speckle noise, erythrocyte spatial distributions, and vis-OCT illumination beam’s incidence angle, may contribute to depth-dependent spectroscopic signal differences in vessels. Therefore, we tuned *z*_*d*_ from 6 *μm* to 14 *μm* in 8 equidistant steps below the identified peak blood signal and computed sO_2_ for each iteration. If a single peak could not be found, perhaps due to noise or spatial averaging, we then tuned *z*_*d*_ from 16 *μm* to 24 *μm* in 8 equidistant steps from the peak amplitude of the anterior vessel wall, which is the typical range where a maximum was visible. The second parameter is the scattering scaling factor *W*, which can vary with erythrocyte spatial distributions and multiple scatterings. Based on *ex-vivo* bovine blood sO_2_ measurements (**Supplementary Materials**) and human retinal sO_2_ measurements, we determined that the strongest regression fits (*R*^2^) were found between *W* = 0.02 and *W* = 0.10. Therefore, we computed 8 iterations of sO_2_ for *W* in this range with a step size of 0.01. The third parameter *S* scales the SDBG amplitude by a small value. Briefly, we measured the SDBG where the vis-OCT signal was attenuated to the noise floor and extrapolated its amplitude at the vessel location by fitting an exponential curve to the SDBG. Due to the low SNR of the measured spectrum relative to the SDBG, small errors in this extrapolation could alter the measured spectrum. To account for these potential errors, we applied a small correction factor *S* to the SDBG as *SB*(*λ, z*) before SDBG subtraction from Eq. 1. We tuned *S* between 0.96 and 1.04 in 9 steps with a step size of 0.01. We measured sO_2_ for each iteration (Fig. 2, Step 8). In total, we calculated sO_2_ for 8 × 8 × 9 = 576 iterations. We stored measured sO_2_ and spectral fit R^2^ for each parameter iteration in 3D matrixes.

### Fine data filtering

To select and remove noise and outliers, we filtered this dataset. First, we rejected all sO_2_ iterations where the spectral fit *R*^2^ < 0.80. We found that noisier arterial spectra resulted in sO_2_ = 100%, or saturation at the maximum possible value. Therefore, we applied stricter rejection criteria for calculations of sO_2_ = 100% and rejected all iterations where *R*^2^ was less than 0.93. Following the data filtering step, we sorted the iterations in ascending order of sO_2_ and selected the 20 central indexes, acting as a pseudo-median measurement. Among these values, we selected the iteration where the *R*^2^ was the highest. We saved the *z*_*n*_, *W*, and *A*_*C*_ for the selected iteration and accepted the sO_2_ with these parameters.

### Vis-OCT systems

We used vis-OCT systems at NYU Langone Health Center (Aurora X1, Opticent Health, Evanston, IL) and Northwestern Medical Hospital (Laboratory Prototype), which were reported, respectively, in our previous work (32). Both systems were fiber-based Michelson interferometers with 30:70 sample to reference splitting ratios. Both systems used telescopic optics in the sample arm, reaching an estimated 7-*μ*m (1/*e*^2^) spot sizes on the retina. The system roll-off is - 4.8 dB/mm in both systems.

### Imaging protocols

Imaging was performed at NYU Langone Health Center and Northwestern Memorial Hospital. We limited light exposure on the cornea to < 250 *μW*, which is considered eye-safe (53). The camera line period was set to 40 *μs* (39 *μs* exposure + 1 *μs* data transfer), or a 25 kHz A-line rate. All volunteers provided informed consent before being imaged. All imaging was approved by respective NYU and Northwestern University Institutional Review Boards adhered to the Tenants of Helsinki. Volunteers had the right to cease imaging without explanation at any stage during imaging.

We performed three scan types for oximetry measurement: (1) arc scan, (2) small FOV raster scan, and (3) large FOV raster scan. An arc scan is a 120-degree segment of a circular scan with a radius of 1.7 mm acquired with 16 B-scans at 8192 A-lines per B-scan. A small-FOV scan is a 1 mm × 1 mm raster scan acquired containing 16 B-scans with 8192 A-lines per B-scan. A large-FOV scan is a 4.8 mm × 4.8 mm raster scan containing 64 B-scans with 4096 A-lines per B-scan. The large-FOV scan was used for the *en-face* oximetry map of the optic disk (Fig. 4). We acquired arc and small-FOV scans in 5s. We acquired the large-FOV scan in 10s. We found no significant differences in sO_2_ values for different scanning modes.

### Vessel selection

We measured sO_2_ in 176 total retinal vessels (98 arteries and 78 veins) across 18 healthy subjects. We analyzed 125 unique retinal vessels (72 arteries and 53 veins) (see **Methods – Retinal oximetry in a healthy cohort**). For vessels with more than one measurement, we selected unique vessels by selecting sO_2_ measurement with the highest R^2^. For repeatability analysis, we selected vessels with at least two repetitions.

### Vessel segmentation

We selected the left and right borders of a vessel, guided by its attenuation shadow. To account for different vessel geometries, we automatically segmented the central 36%, 40%, and 42% of A-lines of the vessel. We repeated these three segmentations for a 4% shift left and right of the detected vessel center, totaling 9 segmentations of the same vessels. We treated each of the 9 segmentations as separate B-scans in the analysis.

### Statistical analysis

To assess whether the lower sO_2_ values were influenced by vessel diameter or were an artifact of poor fitting (lower *R*^2^), we included both parameters in a linear model

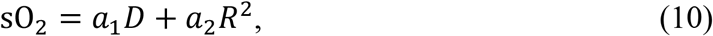

where *D* is vessel diameter; *R*^2^ is the spectral fit R^2^ value; and *a*_1_ and *a*_2_ are arbitrary constants. We did not include the influence of *W* in the above model because we found that it had no significant correlation with vessel diameter, *R*^2^, or sO_2_. A two-way ANOVA was performed in MATLAB 2018a. Significance was considered as *p* < 0.05. We compared sO_2_ measurement populations in Figs. 6b and 6c. We used the two-sample t-test to determine differences in the mean. Significance was considered as *p* < 0.05. t-tests were performed in MATLAB 2018a.

## Acknowledgment

The authors thank Robert Linsenmeier for fruitful discussions and the blood-gas analyzer used in this work. The authors are grateful for the generous support from the National Institutes of Health (R01EY019949, R01EY026078, R01EY028304, R01EY029121, R44EY026466, and T32EY25202).

## Disclosure

HFZ, RVK, and YW have financial interests in Opticent Health

## Supplementary information

### *Ex vivo* phantom verification and comparison

**Figure S1.**
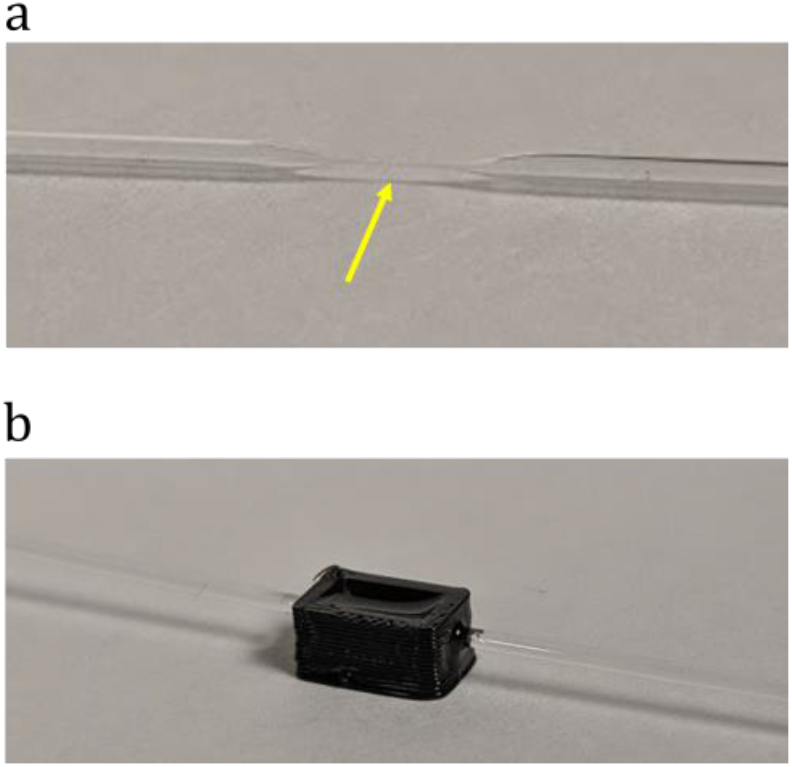
Capillary tube phantom for *ex-vivo* vis-OCT oximetry. (a) Glass capillary tube pulled to an inner diameter of ∼ 200 *μ*m (arrow); (b) Tube inserted in a homemade well under ∼ 500 *μ*m of immersion oil.

We measured sO_2_ values in an *ex vivo* bovine blood phantom using vis-OCT and compared them with a blood-gas analyzer (Rapidlab 248, Siemens Healthcare Diagnostics, Malvern, PA). Briefly, we pulled a glass capillary tube to an inner diameter of ∼ 200 *μ*m (Fig. S1a). We embedded the pulled tube in the middle of a homemade plastic well (Fig. S1b). To reduce specular reflections from the air-glass interface, we added immersion oil (refractive index = 1.52) to the well until the tube was covered by ∼ 500 *μ*m of oil.

Next, we prepared oxygenated (sO_2_ ≈ 100%) and deoxygenated (sO_2_ ≈ 0%) bovine blood (Quadfive, Ryegate, MT). Hematocrit of blood samples was 45%. To oxygenate blood, we exposed it to a constant stream of oxygen while mixing with a magnetic stir bar. We verified that the blood was oxygenated using the blood-gas machine. To deoxygenate blood, we added sodium dithionite to the solution (54). We monitored the partial pressure of oxygen (pO_2_), partial pressure of carbon dioxide (pCO_2_), pH, and temperature of the mixture using the blood-gas machine and converted to sO_2_ (55). We continued adding sodium dithionite and measuring sO_2_ until the blood was sufficiently deoxygenated. Following oxygenation and deoxygenation, we immediately loaded blood samples into syringes to prevent influences from ambient air.

We used oxygenated and deoxygenated samples to make 17 blood samples between sO_2_ ≈ 100% and sO_2_ ≈ 0%. To this end, we mixed oxygenated and deoxygenated samples to create blood of another oxygenation level. We measured sO_2_ of the mixed blood using the blood-gas machine. We imaged each blood sample immediately after blood-gas machine measurement.

Before loading blood into the tube, we flushed the tube with a phosphate-buffered saline (PBS) and heparin solution to prevent clotting or sedimentation. Then, we loaded the blood into a syringe, which was connected to the glass tube by ∼ 1 m of plastic tubing. We placed the syringe in a syringe pump, which flowed the blood at ∼ 0.03 mm/s inside the glass tube to prevent clotting or sedimentation.

Before imaging, we focused the beam on the tube by adjusting tube height and maximizing the intensity of backscattered light. After reaching best focus, we adjusted the reference arm to place the top of the tube < 100 *μ*m from the zero-delay. Then, we imaged the tube using a 512 × 512 raster scan. Optical power incident on the tube was 1.20 mW. After imaging each blood sample, we re-flushed the tube with the PBS and heparin solution.

We measured sO_2_ with vis-OCT in each blood sample using the AS-OCT processing proposed in this work. Since scattering factor *W* was not well-agreed upon in the literature, we varied *W* to find the highest spectral fit R^2^. We found that best spectral fit R^2^ was reached between 0.02 ≤ *W* ≤ 0.10.

We computed 100 vis-OCT sO_2_ measurements for each tube. Briefly, we processed and stored all 512 B-scans. Then, we randomly selected and averaged 50 different B-scans from this set for sO_2_ measurement. Then, we refreshed the 512 B-scans and repeated random selection 100 times to reach 100 sO_2_ measurements.

**Figure S2.**
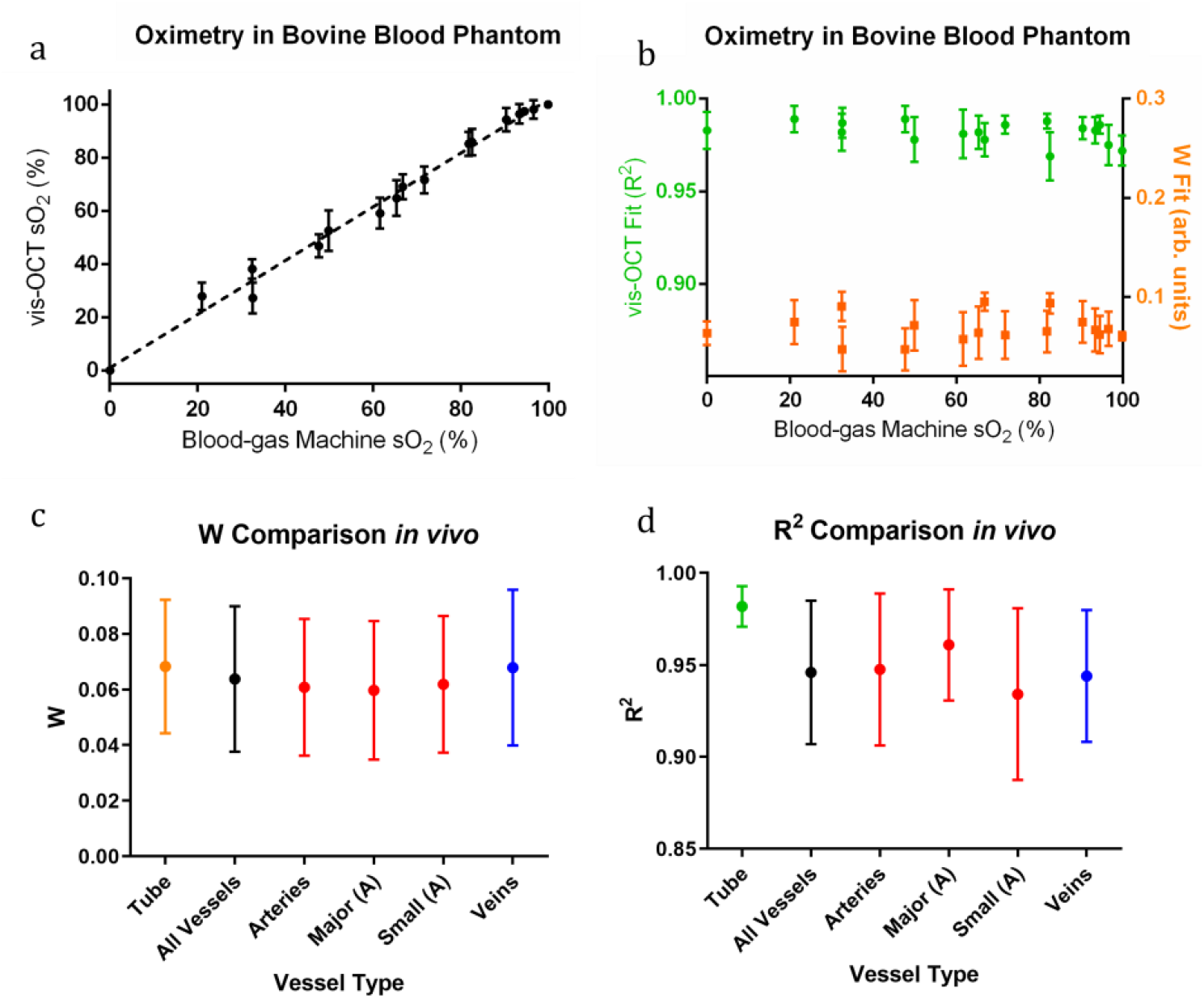
Results of vis-OCT oximetry in *ex vivo* phantom and comparison to *in vivo* human eye. (a) vis-OCT sO_2_ measurements in phantom plotted against sO_2_ measurements from blood-gas machine; (b) Distributions of spectral fit R^2^ and best fit *W* in phantom; (c) Distributions of best fit *W* in phantom compared to human eye; (d) Distributions of best fit R^2^ in phantom compared to human eye

Fig. S2a shows tube sO_2_ measured by vis-OCT and the blood-gas machine for 17 tubes ranging from sO_2_ ≈ 0% to sO_2_ ≈ 100%. The equation of the best fit line is *y* = 1.01*x* + 1.28 and the coefficient of regression is *R*^2^ = 0.97. The relationship between the blood-gas machine and vis-OCT sO_2_ was nearly a slope of 1 with only ∼1% bias. This was within our target accuracy, so we did not apply a post-hoc calibration curve to vis-OCT measurements in this work.

Fig. S2b shows spectral fits (R^2^) and best fit *W* for each tube. Average spectral fit R^2^ was 0.98 and average W was 0.068. In Fig. S2c, we plot the best-fit *W ex vivo* and *in vivo*. In the tubes, we measured *W* = 0.068 ± 0.024. For all unique vessels, we measured *W* = 0.064 ± 0.026, agreeing well with the tube data. W measurements were 0.061 ± 0.025, 0.060 ±0.025, 0.062 ± 0.025, and 0.068 ± 0.028 for all arteries, large arteries, small arteries, and all veins, respectively. In Fig. S2d, we plot the spectral fit R^2^ *ex vivo* and *in vivo*. In tubes, we measured R^2^ = 0.98 ± 0.01. For all unique vessels, we measured average R^2^ = 0.95 ± 0.04. R^2^ measurements were 0.95 ± 0.04, 0.96 ±0.03, 0.93 ± 0.05, and 0.94 ± 0.04 for all arteries, major (diameter ≥ 100 *μ*m) arteries, small arteries (diameter < 100 *μ*m), and all veins, respectively. We anticipated that major arteries would have slightly higher R^2^ (0.96) than smaller ones (0.93), considering the increased spatial averaging for larger vessels. Nevertheless, R^2^ did not have a significant influence on sO_2_ value (see **Retinal oximetry in a healthy cohort**).

**Table S1.** All unique vessels analyzed for repeatability in Fig. 4c. ‘A’ indicates artery and ‘V’ indicates vein.

| Unique Vessel | Type | Avg. Diameter ($\mu\text{m}$) | Avg. $\text{sO}_2$ (%) | Repetitions | $\text{sO}_2$ Standard Deviation (%) | Avg. Spectral Fit $R^2$ |
| --- | --- | --- | --- | --- | --- | --- |
| 1 | A | 138 | 100.0 | 3 | 0 | 0.972 |
| 2 | A | 134 | 98.9 | 2 | 1.63 | 0.980 |
| 3 | A | 130 | 100.0 | 2 | 0 | 0.963 |
| 4 | A | 128 | 97.1 | 2 | 1.06 | 0.961 |
| 5 | A | 124 | 99.6 | 2 | 0.64 | 0.965 |
| 6 | A | 122 | 96.2 | 3 | 3.75 | 0.971 |
| 7 | A | 122 | 97.8 | 4 | 3.81 | 0.962 |
| 8 | A | 121 | 99.2 | 2 | 1.20 | 0.946 |
| 9 | A | 121 | 99.9 | 3 | 0.23 | 0.969 |
| 10 | A | 120 | 100.0 | 2 | 0 | 0.955 |
| 11 | A | 119 | 95.5 | 2 | 4.88 | 0.948 |
| 12 | A | 116 | 99.6 | 2 | 0.64 | 0.953 |
| 13 | A | 116 | 98.4 | 2 | 2.26 | 0.970 |
| 14 | A | 116 | 98.6 | 3 | 2.48 | 0.943 |
| 15 | A | 113 | 97.0 | 2 | 4.24 | 0.966 |
| 16 | A | 105 | 96.1 | 2 | 5.59 | 0.948 |
| 17 | A | 104 | 97.9 | 2 | 2.97 | 0.964 |
| 18 | A | 93 | 92.0 | 2 | 2.83 | 0.966 |
| 19 | A | 89 | 99.6 | 2 | 0.64 | 0.963 |
| 20 | A | 89 | 96.2 | 2 | 1.27 | 0.932 |
| 21 | A | 86 | 91.1 | 2 | 2.40 | 0.950 |
| 22 | A | 86 | 92.0 | 3 | 6.83 | 0.918 |
| 23 | A | 62 | 92.9 | 2 | 1.13 | 0.958 |
| 24 | A | 48 | 91.3 | 2 | 2.55 | 0.912 |
| 25 | V | 174 | 62.3 | 2 | 3.46 | 0.968 |
| 26 | V | 160 | 57.0 | 2 | 2.12 | 0.953 |
| 27 | V | 160 | 58.1 | 2 | 0.35 | 0.954 |
| 28 | V | 157 | 54.8 | 3 | 3.07 | 0.943 |
| 29 | V | 156 | 60 | 3 | 2.31 | 0.967 |
| 30 | V | 153 | 55.4 | 3 | 2.97 | 0.957 |
| 31 | V | 149 | 55.9 | 4 | 4.98 | 0.951 |
| 32 | V | 146 | 56.0 | 2 | 1.41 | 0.954 |
| 33 | V | 145 | 61.0 | 3 | 2.06 | 0.927 |
| 34 | V | 139 | 58.4 | 3 | 3.44 | 0.918 |
| 35 | V | 135 | 55.0 | 2 | 1.20 | 0.900 |
| 36 | V | 133 | 60.1 | 2 | 1.27 | 0.974 |
| 37 | V | 131 | 54.3 | 2 | 0.49 | 0.940 |
| 38 | V | 128 | 51.5 | 2 | 4.45 | 0.955 |
| 39 | V | 125 | 50.6 | 2 | 0.78 | 0.980 |
| 40 | V | 80 | 59.0 | 2 | 2.4 | 0.977 |
| 41 | V | 70 | 64.1 | 2 | 3.46 | 0.913 |
| 42 | V | 62 | 57.7 | 2 | 1.41 | 0.904 |

### Comparing depth-averaging with slope for estimation of attenuation coefficient

**Table S2.**
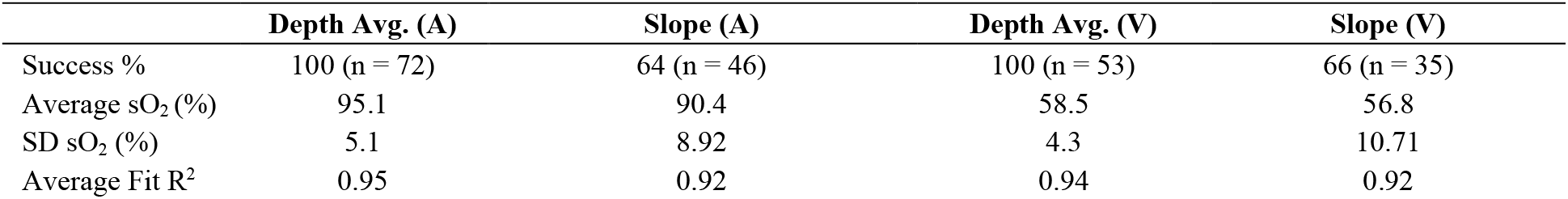
Comparison of depth-averaging and slope methods for vis-OCT retinal oximetry

**Figure S3.**
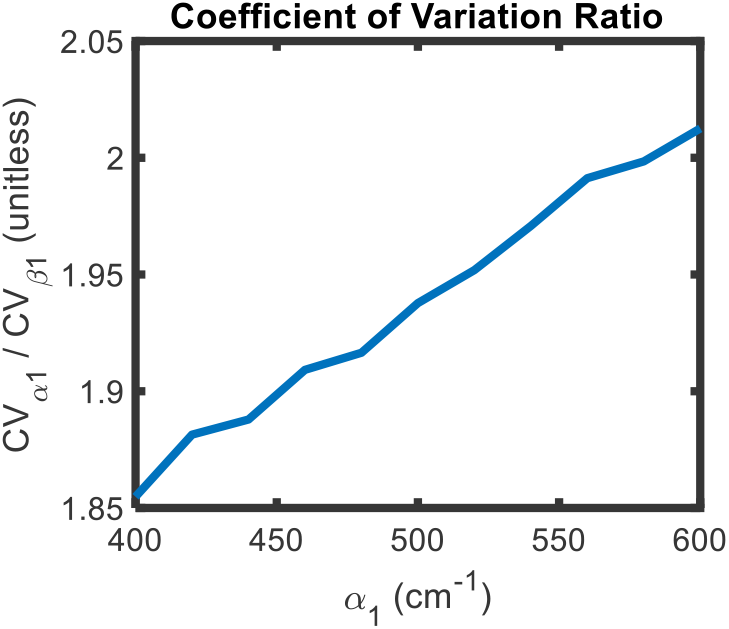
Coefficient of variation ratio between slope and depth averaging methods for estimating attenuation coefficient. Exponential decay model from Eq. S3 is used for calculations. *α*_1_ is attenuation coefficient estimated by slope method; *β*_1_ is proportional to *α*_1_ and estimated by depth averaging method.

Empirically, we found that depth-averaging the natural logarithm of the SDA-lines yielded less noisy spectra, as compared with the slope method (**see Methods – Comparison of depth-averaging and slope methods**). We verified these empirical observations by Monte Carlo simulation. To begin, we applied the slope method and depth-averaging method to the equation of a line, which is predicted by Eqn. 7 in **Methods**. (for simplification, removing small effect of LCA):

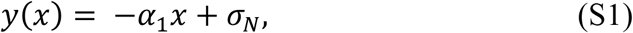

where *α*_1_ is an arbitrary constant and *σ*_*N*_ is random, normally distributed noise. *x* was a 30-pixel vector ranging from 0 to 35 *μm* for the depth selection window. We used the slope method to compute the slope of Eq. S1 and directly find *α*_1_. We used the depth-averaging method to compute the average value of Eq. S1 (similar to Eq. 8) to find the constant *β*_1_ *∝ α*_1_. We computed 10^5^ iterations of such measurements, and then measured their coefficients of variation:

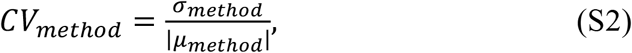

where *CV*_*method*_ is the coefficient of variation of the measured *α*_1_ or *β*_1_ from each respective method, *σ*_*method*_ is the SD of the measured *α*_1_ or *β*_1_ from each respective method, and *μ*_*met*h*od*_ is the average of the measured *α*_1_ or *β*_1_ from each respective method. We computed *CV*_*method*_ for *α*_1_ = 400 cm^-1^ to *α*_1_ = 600 cm^-1^ in an increment of 20 cm^-1^, which covered the reported attenuation coefficients of blood in the visible-light spectral range for *W* = 0.064. We used normally distributed noise with amplitude *σ*_*N*_ = 0.02 (arb. units), which was a relative noise typically observed *in vivo*. For each value of *α*_1_, we calculated 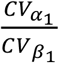, which compared the relative uncertainty of the slope method to the depth-averaging method. For all values of *α*, 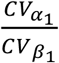 converged to 1.67. In general, we found that value of 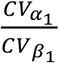 was independent of *α*_1_and *σ*_*N*_. This suggested that the depth-averaging method had an intrinsic noise reduction advantage of 67% over the slope method for additive, normally distributed noise.

In reality, however, the SDA-lines follow an exponential decay with the additive noise. We applied a natural logarithm to this function:

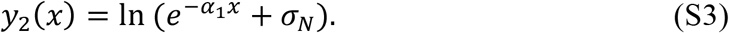

One frequent assumption by the slope method is that signal is significantly greater than noise, or 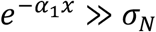, after which Eq. S3 would converge to a noiseless version of Eq. S1. However, in the human retina, SNR is often low, and this assumption might not be correct. To this end, the noise in Eq. S3 after the natural logarithm is less trivial than in Eq. S1 since there is no longer a linear relationship between *α*_1_ and *σ*_*N*_. We repeated the simulation described above, except we generated the signal and noise using Eq. S3. Fig. S3 plots 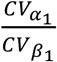 for this simulation. Like the analysis from Eq. S1, the coefficient of variation using the slope method is always higher than that using the depth averaging method. However, the 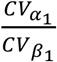 is not constant and increases with increased *α*_1_. This is because the relationship between *α*_1_ and *σ*_*N*_ is nonlinear in Eq. S3. This has important implications for the slope method in sO_2_ calculation, since the measured blood spectrum can have different noise levels for different wavelengths and for different depth selection windows. In this work, we demonstrated empirically that depth averaging is statistically advantageous over the slope method for sO_2_ calculation, consistent with the simulation.

### Longitudinal Chromatic Aberration vis-OCT Retinal Oximetry

**Figure S4.**
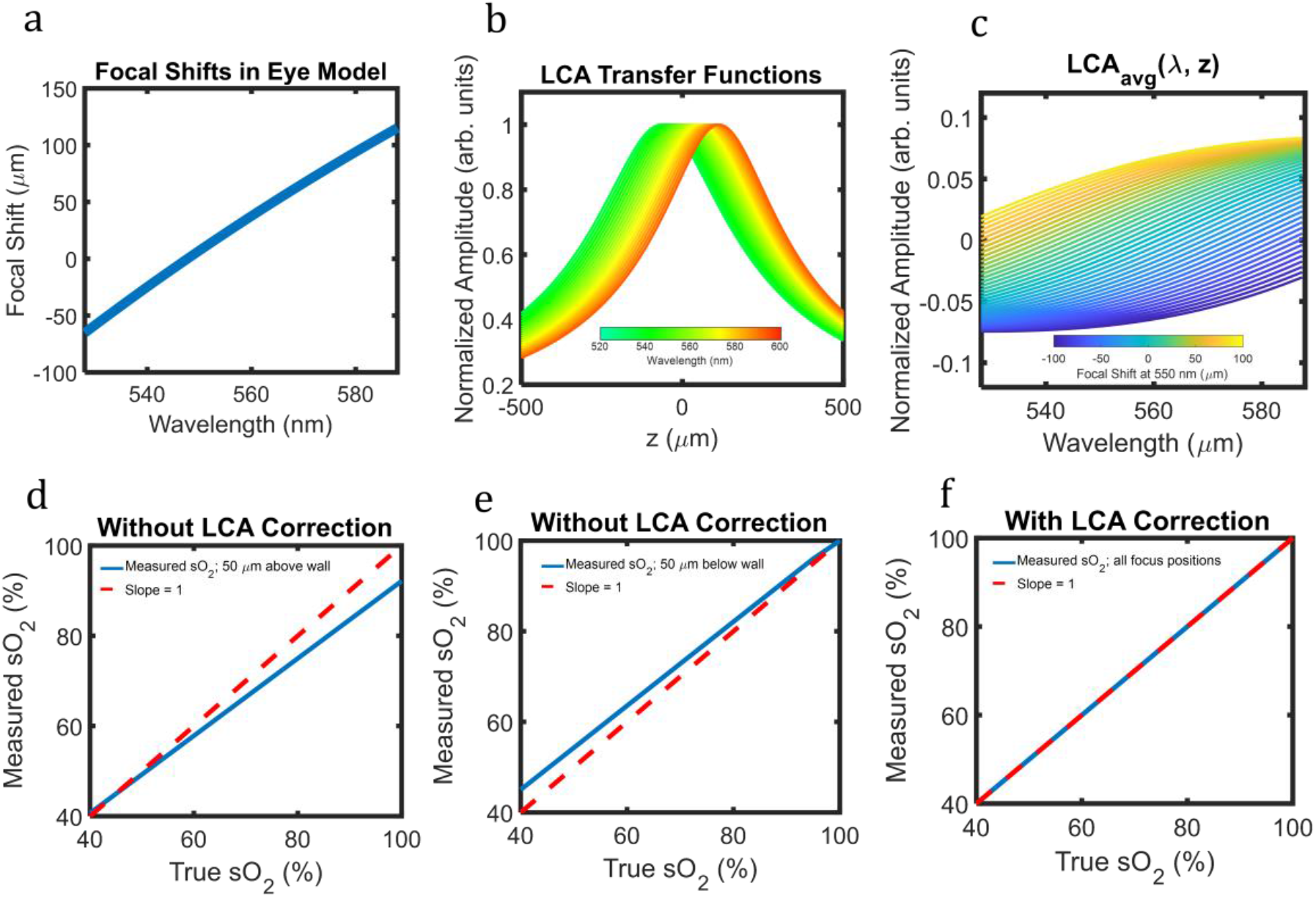
Simulation of LCA in human eye and influence on vis-OCT retinal oximetry. (a) CFS in human eye simulated by Zemax software; (b) Transfer function of the LCA on vis-OCT SDA-lines. Colors depict central wavelength of STFT window; (c) Simulated LCA contribution to measured spectrum after AS-OCT processing; (d) Simulated sO_2_ measurement without LCA correction when focus at 550 nm is 50 *μ*m above the anterior vessel wall; (e) Simulated sO_2_ measurement without LCA correction when focus at 550 nm is 50 *μ*m below the anterior vessel wall; (f) Simulated sO_2_ measurement with LCA correction for all focus positions

We developed an approach for fitting LCA transfer functions to sO_2_ measurement using the physical optics of the human eye. First, we simulated the CFS in the human eye model from Polans et al. (56) using Optic Studio 16 (Zemax, Kirkland, Washington). Since the wavelength ranges and lateral resolutions of the vis-OCT systems used in this study were approximately the same (see **Methods – Vis-OCT Systems**), we used the same CFS for both systems (Fig. S4a). The simulated chromatic focal shift (CFS) CFS is consistent with that previously measured in the human eye (57). Then, we calculated potential LCA transfer functions using a modified version of the equation used in (58) to account for spectroscopic analysis:

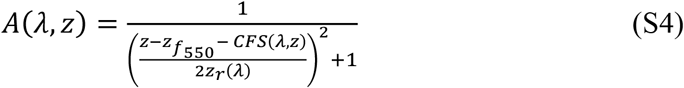

where 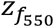 is the reference focusing depth at 550 nm, *CFS*(*λ, z*) are the chromatic focal shifts, and *z*_*r*_(*λ*) are the wavelength-dependent Rayleigh lengths (assumed refractive index = 1.35 and 1/*e*^2^ spot size = 7 *μ*m). We calculated 41 LCA transfer functions up to 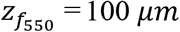 above and below the anterior wall of a simulated vessel in 5 *μm* increments. Fig. S4b illustrates a simulated LCA transfer function 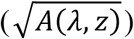 for 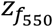 focused at the anterior vessel wall. Then, we normalized 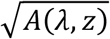 by its spectral profile at z_*n*_ = 12 *μ*m into the simulated vessel lumen, consistent with typical sO_2_ measurements, and took its natural logarithm. We found *LCA*_*avg*_(*λ, z*) by averaging the normalized ln 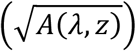 from *z*_0_ = 16 *μ*m to *z*_0_ + Δ*z* = 46 *μ*m into the vessel lumen (Eqn. 7), also consistent with typical sO_2_ measurements. To create the LCA lookup table, we saved *LCA*_*avg*_(*λ, z*) for each of the 41 focal positions (Fig. S4c).

To understand potential influence of LCA on sO_2_ measurement, we simulated SDA-lines in a vessel consistent with the Beer-Lambert law and the attenuation spectra in Faber et. al (23). We multiplied 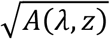 at each focal position with the SDA-lines to account for LCA (Eqn. 1) and took its natural logarithm. We averaged the spectrum at the same depths used to find *LCA*_*avg*_(*λ, z*) (Eqn. 8 in **Methods**). We noted that for all simulated physiological sO_2_ measurements (sO_2_ = 40% to sO_2_ = 100%) and focal positions, the peak-to-peak amplitude of *LCA*_*avg*_(*λ, z*) was less than 0.25 times the peak-to-peak amplitude of 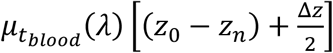. We used this relationship to constrain physically reasonable *LCA*_*avg*_(*λ, z*) to avoid overfitting this parameter in the sO_2_ measurement. Furthermore, since the above constraint described relative amplitudes only, it was independent optical power incident on the vessel.

We measured sO_2_ in the above simulation without and with LCA fitting described in **Methods – Oximetry Fitting Model**. We measured sO_2_ for up to 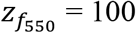 *μm* above and below the anterior wall of a simulated vessel in 5 *μm* increments. Measurements were derived from simulated SDA-lines of oxygen-dependent spectra from sO_2_ = 40% to 100%. Figs. S4d and S4e plot sO_2_ measurements without fitting the contribution of *LCA*_*avg*_(*λ, z*). Fig. S4d shows sO_2_ measurements for 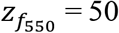 *μm* above the vessel anterior wall and Fig. S4e shows sO_2_ measurements for 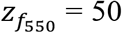 *μm* below the anterior wall. When 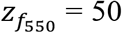 *μm* above the anterior wall, sO_2_ is underestimated. When 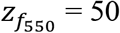 *μm* below the anterior wall, sO_2_ is overestimated. Fig. S4f plots sO_2_ measurements with fitting the contribution of *LCA*_*avg*_(*λ, z*), as described in **Methods – Oximetry Fitting Model**. For all values of 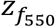, measured sO_2_ matches the ground truth sO_2_ with a slope of 1.

We recognize this is a simplified approach may not have fully appreciated the exact influence of LCA in each recorded image. Nevertheless, such corrections are based on the well-verified aberrations and defocusing in the human eye. More precise LCA correction may be reached with a wavefront sensor and adaptive optics to directly measure wavelength-dependent aberrations, although they add expense and complexity to the vis-OCT system. The influence of LCA can also be reduced by employing an achromatizing lens (30) in the sample arm of the system. Additionally, our vis-OCT systems used a focusing beam diameter (1/*e*^2^) of 7 *μm*. Decreasing beam diameter at the cornea and increasing depth of focus can also reduce influence of LCA on sO_2_ calculation, although it may challenge laterally resolving smaller vessels.

### Depth-resolved spectral analysis

**Figure S5.**
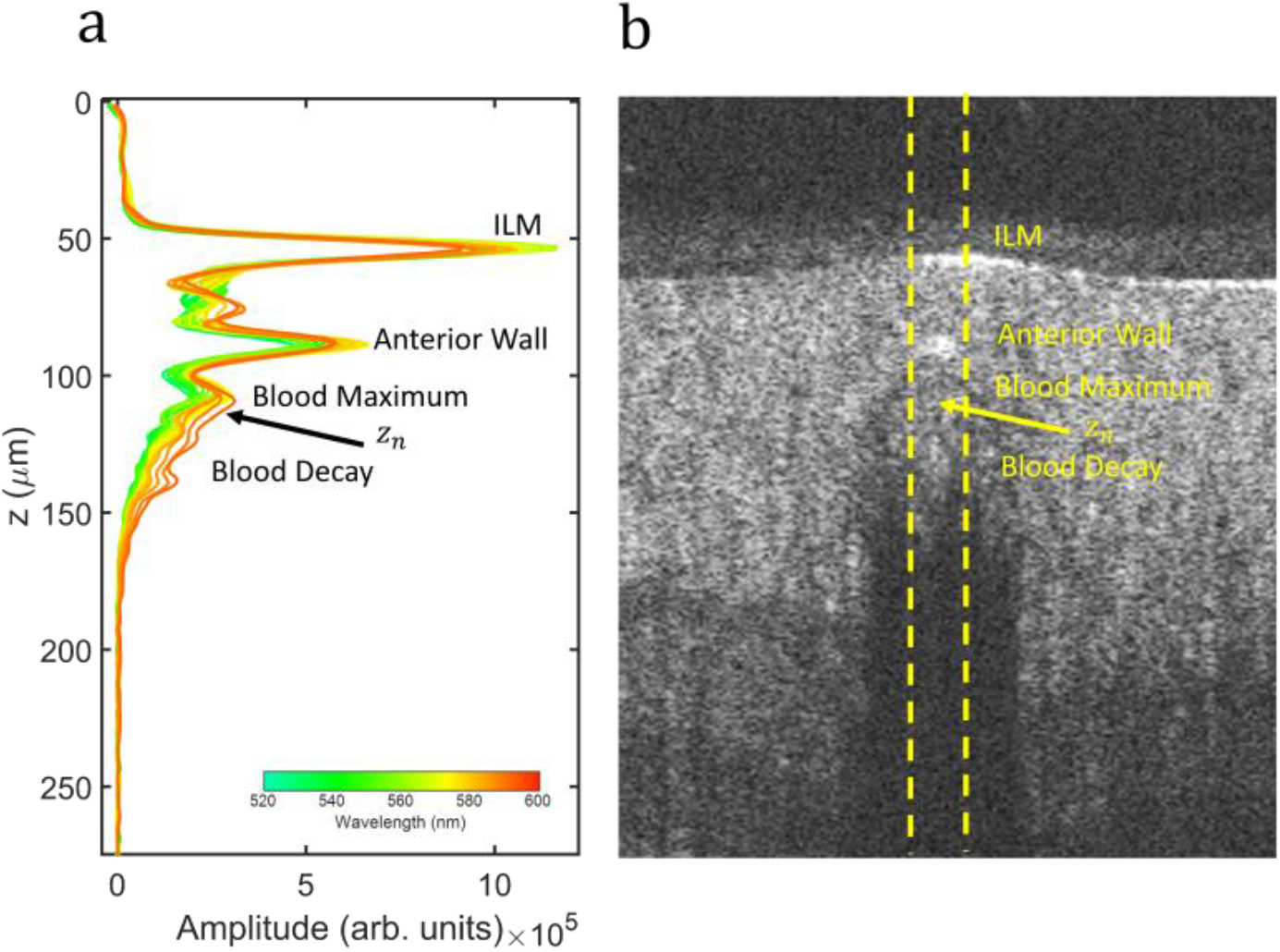
SDA-lines features. (a) SDA-lines from vein in human retina. Color bar represents central wavelength of STFT window. *z*_*n*_ indicates depth of normalization where SDA-lines start to decay in amplitude; (b) Magnified B-scan where SDA-lines in were measured. SDA-lines were averaged laterally within yellow dashed lines

**Figure S6.**
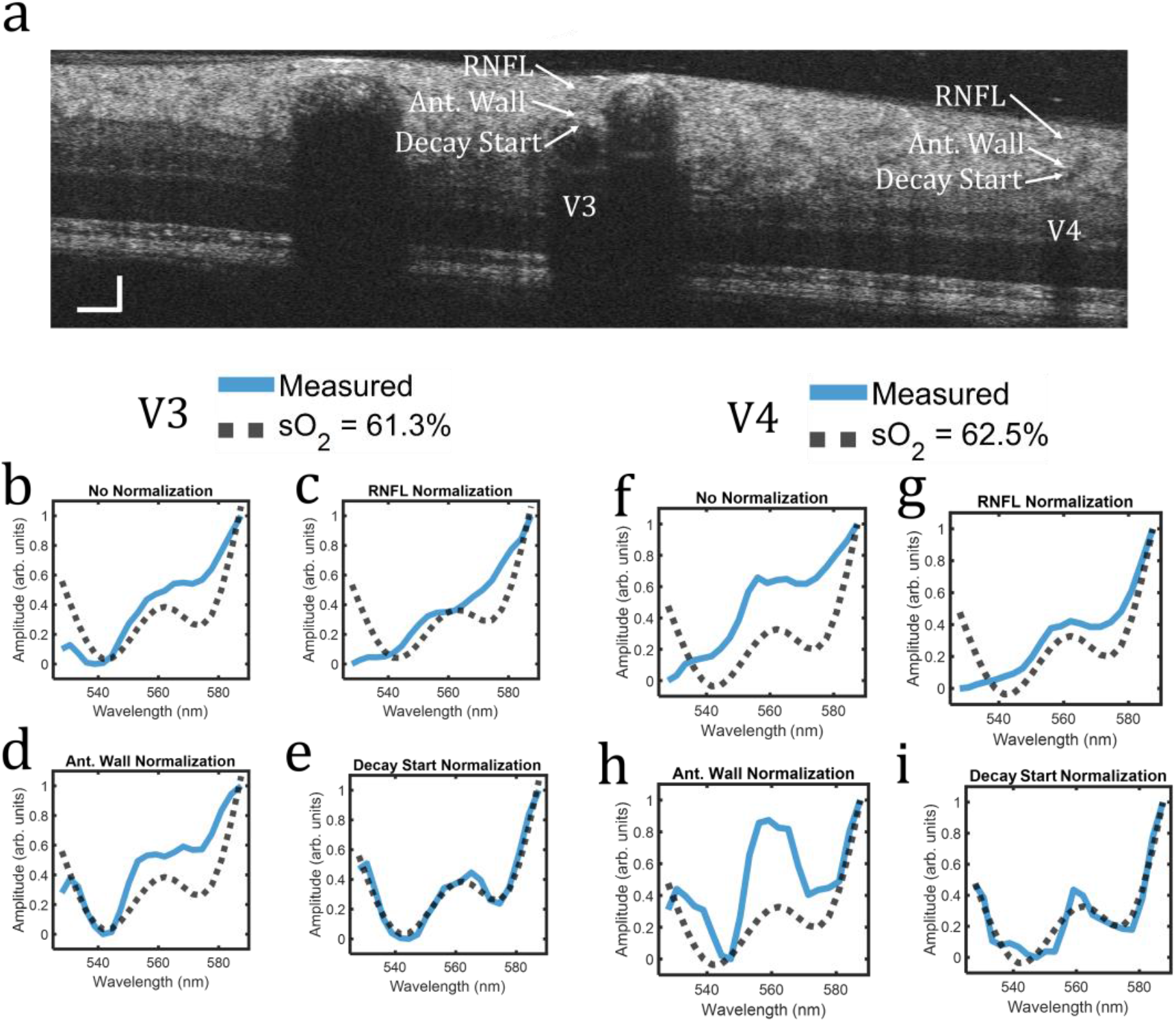
Spectroscopic normalizations in the human retina. (a) vis-OCT B-scan. Vessels labeled V3 and V4 both identified as veins. Arrows highlight anatomical features used for normalization. (b-e) Measured spectrum (blue line) and literature-derived spectrum for sO_2_ = 61.3% (back dashed line) in V3 for no normalization, normalization by the RNFL, normalization by the vessel anterior wall, and normalization by the start of signal decay in blood, respectively. (f-i) The same analysis for Fig. S6b-S6e is replicated for V4.

Fig. S6a shows the same vis-OCT B-scan as in Fig. 3a (see **Methods**), with vessels V3 and V4, two veins, highlighted. We tested spectroscopic normalizations in V3 and V4 (Fig. S6b-S6i) in the same ways depicted in Figs. 3b-3i. Like in Figs. 3b-3i, the only normalization agreeing well with the literature is the decay start normalization, indicating removal of spectral contaminants. The other normalizations do not agree well with the literature and are not necessarily consistent across different vessels. For example, the anterior wall normalization for V3 (Fig. S6d) shows agreement between the measured and predicted spectrum only for wavelengths shorter than 540 nm. Such trend is not seen in V4 (Fig. S6h), where the measured spectrum has a higher contrast “W” shape. Notably, vessels V3 and V4 are smaller in diameter and buried deeper under the RNFL than vessels V1 and V2 in Fig. 3. Nevertheless, removal of spectral contaminants allows for accurate spectral measurement.

**Figure S7.**
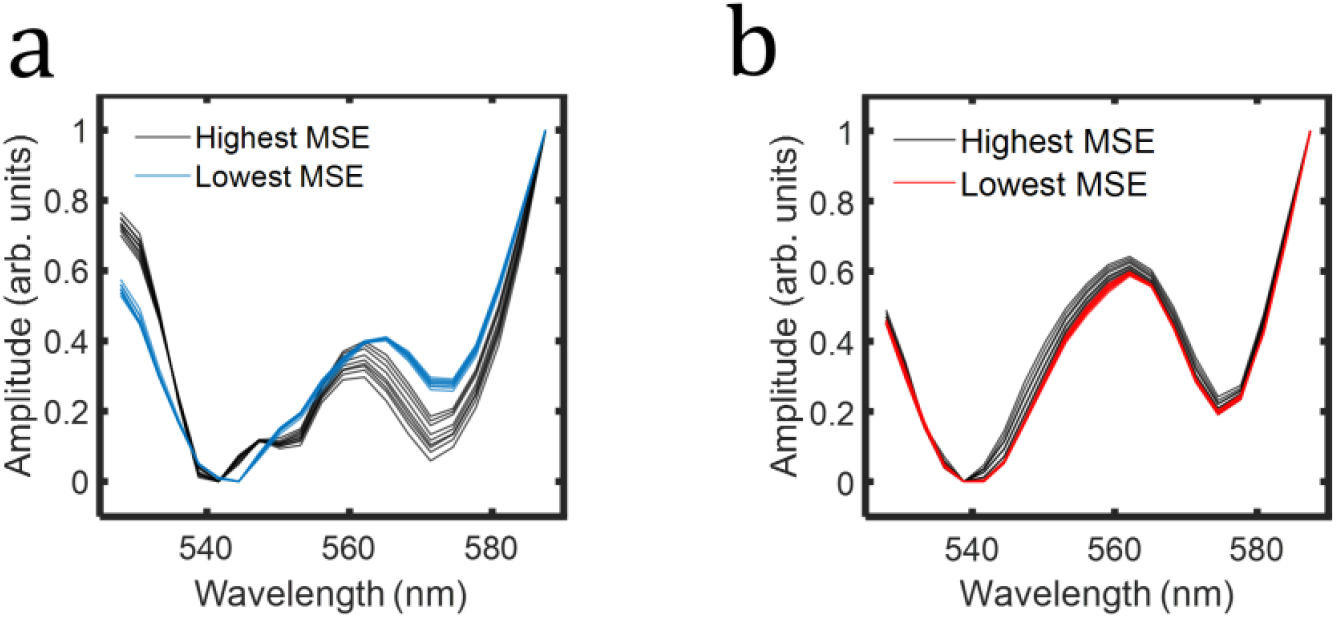
Measured STFT spectra from the highest and lowest mean-squared-error (MSE) in the spectral stability matrixes in Fig. 4. (a) Measured STFT spectra from Fig. 4a. Black lines plot nine spectra after depth perturbations for the highest MSE (black box in Fig. 4a) and blue lines plot nine spectra after depth perturbations for the lowest MSE (green box in Fig. 4a). (b) Same analysis as (a) but for the spectral stability matrix in Fig. 4b. Red lines indicate lowest MSE.

**Figure S8.**
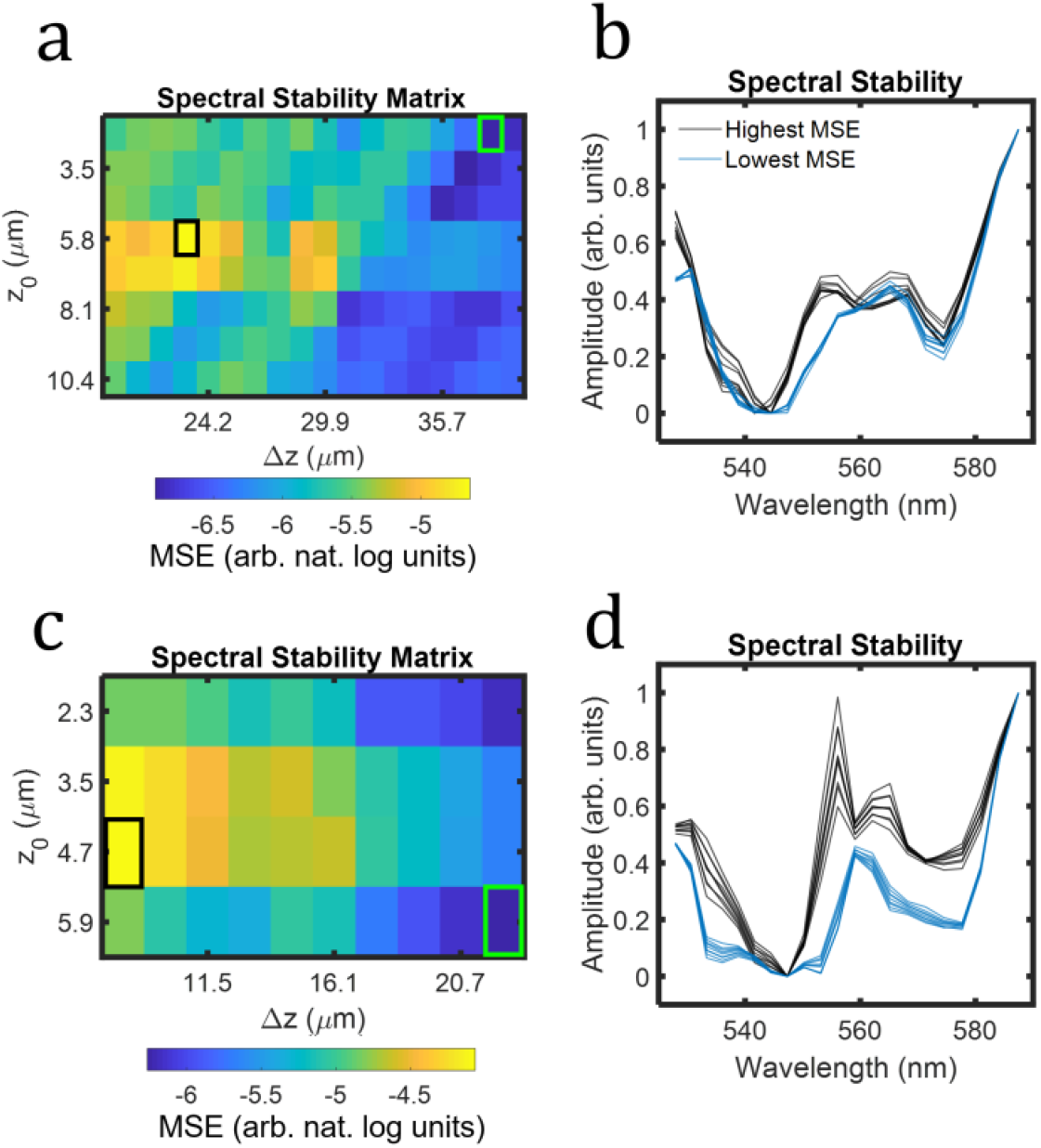
Spectral stability analysis in human retinal vessels. (a) Spectral stability matrix (SSM) for vessel V3 from Fig. S6a. Green box highlights lowest mean-squared-error (MSE) and black box highlights highest MSE; (b) Spectra in V3 after nine depth perturbations for the lowest MSE (blue lines) and highest MSE (black lines) in Fig. S8a, respectively. (c) SSM for vessel V4 from Fig. S6a. Green box highlights MSE and black box highlights highest MSE. (d) Spectra in V4 after nine depth perturbations for the lowest MSE (red lines) and highest MSE (black lines) in Fig. S8c, respectively.

Fig. S8 shows the spectral stability analysis for vessels V3 and V4 in Fig. S6a. The analysis is the same as that shown for vessels V1 and V2 in Figs. 4 & S7. The spectra selected at the lowest MSE of the SSM (Figs S8b and S8d) are stable in response to depth perturbations and are therefore consistent with the Beer-Lambert model of attenuation. We note that Fig. S8c shows a smaller depth range than Fig. S7a, which is due to the smaller profile of the vessel.

## Notes

### Competing Interest Statement

HFZ, RVK, and YW have financial interest in Opticent Health

